# CHCHD2 mutant mice display mitochondrial protein accumulation and disrupted energy metabolism

**DOI:** 10.1101/2024.08.30.610586

**Authors:** Szu-Chi Liao, Kohei Kano, Sadhna Phanse, Mai Nguyen, Elyssa Margolis, YuHong Fu, Jonathan Meng, Mohamed Taha Moutaoufik, Zac Chatterton, Hiroyuki Aoki, Jeffrey Simms, Ivy Hsieh, Felecia Suteja, Yoshitaka Sei, Eric J. Huang, Kevin McAvoy, Giovanni Manfredi, Glenda Halliday, Mohan Babu, Ken Nakamura

**Affiliations:** Gladstone Institute of Neurological Disease, Gladstone Institutes, San Francisco, CA; Department of Nutritional Sciences & Toxicology, University of California Berkeley, Berkeley, CA; Endocrinology Graduate Program, University of California Berkeley, Berkeley, CA; Aligning Science Across Parkinson’s (ASAP) Collaborative Research Network, Chevy Chase, MD; Department of Biochemistry, University of Regina, Regina, Saskatchewan, Canada; Weill Institute for Neurosciences, Department of Neurology, University of California San Francisco, San Francisco, CA; Neuroscience Graduate Program, University of California San Francisco, San Francisco, CA; Brain and Mind Centre & Faculty of Medicine and Health School of Medical Sciences, University of Sydney, Sydney, Australia; Department of Pathology, University of California San Francisco, San Francisco, CA; Biomedical Sciences Graduate Program, University of California San Francisco, San Francisco, CA; Feil Family Brain and Mind Research Institute, Weill Cornell Medicine, New York, NY

## Abstract

Mutations in the mitochondrial cristae protein CHCHD2 lead to a late-onset autosomal dominant form of Parkinson’s disease (PD) which closely resembles idiopathic PD, providing the opportunity to gain new insights into the mechanisms of mitochondrial dysfunction contributing to PD. To begin to address this, we used CRISPR genome-editing to generate CHCHD2 T61I point mutant mice. CHCHD2 T61I mice had normal viability, and had only subtle motor deficits with no signs of premature dopaminergic (DA) neuron degeneration. Nonetheless, CHCHD2 T61I mice exhibited robust molecular changes in the brain including increased CHCHD2 insolubility, accumulation of CHCHD2 protein preferentially in the substantia nigra (SN), and elevated levels of α-synuclein. Metabolic analyses revealed an increase in glucose metabolism through glycolysis relative to the TCA cycle with increased respiratory exchange ratio, and immune-electron microscopy revelated disrupted mitochondria in DA neurons. Moreover, spatial genomics revealed decreased expression of mitochondrial complex I and III respiratory chain proteins, while proteomics revealed increased respiratory chain and other mitochondrial protein-protein interactions. As such, the CHCHD2 T61I point-mutation mice exhibit robust mitochondrial disruption and a consequent metabolic shift towards glycolysis. These findings thus establish CHCHD2 T61I mice as a new model for mitochondrial-based PD, and implicate disrupted respiratory chain function as a likely causative driver.

## INTRODUCTION

Mitochondria have long been hypothesized to play a central role in the pathophysiology of Parkinson’s disease (PD). Indeed, sporadic PD has been repeatedly associated with decreased activity of respiratory chain function, especially mitochondrial complex I in the substantia nigra pars compacta (SNc) (*1–3*). Aging is also the greatest risk factor for PD, and mitochondrial DNA mutations accumulate at a faster rate in SNc dopaminergic (DA) neurons in PD than in normal aging (*4*). Moreover, neurotoxin data suggest that SNc DA neurons are preferentially vulnerable to mitochondrial complex I dysfunction, as exposure to the mitochondrial complex I inhibitors rotenone and 1-methyl-4-phenyl-1,2,3,6-tetrahydropyridine (MPTP) either predispose to PD and increase the risk of (rotenone) or cause a parkinsonian syndrome (MPTP) (*5–7*).

Although the above neuropathologic and neurotoxin data implicate mitochondria in PD pathophysiology, they do not establish that changes in mitochondria actually cause PD. As such the discoveries that mutations in the mitochondrial protein PINK1 and the mitochondria-associated protein Parkin cause familial forms of PD (*8*) are particularly important in proving that mitochondrial dysfunction can cause PD. Moreover, the roles of PINK1 and Parkin in promoting mitochondrial turnover, mitochondrial respiration and bioenergetics reveal potential mechanisms by which mitochondria may cause PD (*9–13*). Nonetheless, most patients with *Parkin* mutations, the most frequent cause of autosomal recessive PD, do not have Lewy body pathology (*14*) leading many investigators to hypothesize that autosomal recessive forms of PD are distinct disease subtypes from autosomal dominantly-inherited and sporadic forms of PD that more closely resemble each other (*15*).

As such, the discovery that mutations in the mitochondrial cristae protein CHCHD2 (coiled-coil-helix-coiled-coil-helix domain containing 2) cause an autosomal dominant form of late-onset PD that closely resembles sporadic PD (*16*), including widespread synuclein deposition (*17*), provides a potentially critical link proving that mitochondrial dysfunction can indeed cause a late onset form of PD that is typical of both sporadic PD and autosomal dominant PD. It also provides an important opportunity to learn how mitochondrial dysfunction can cause PD. Understanding how mutations in CHCHD2 promote PD has proven challenging, however. Although CHCHD2 may in some way support respiration (*18, 19, 20*), its physiologic function(s) remain unknown, especially in vivo. Moreover, mice lacking CHCHD2 have only subtle phenotypes (*21, 22*), suggesting that CHCHD2 mutations may produce toxicity through gain of a toxic function. Indeed, mutations in CHCHD10, CHCHD2’s paralog and binding partner, appears to produce toxicity through gain of function, suggesting that CHCHD2 may have similar effects (*23*), a possibility also supported by recently described mouse models with mutant CHCHD2 (*24, 25*).

However, a significant challenge in PD research remains the lack of well characterized mouse models for monogenic PD that exhibit dysfunction and preferential degeneration in nigrostriatal dopamine neurons, the pathological hallmark of the disease.

## RESULTS

### CHCHD2 T61I mice generation and genotyping

CHCHD2 p.T61I mice were made with CRISPR/Cas9 genome-editing (Fig. 1A). Specifically, two sgRNAs were used to remove the entire exon 2 and exon 3, and a targeting donor vector was used to deliver the cytosine-to-thymine mutation, resulting in the codon change from threonine to isoleucine. This approach allowed us to specifically target *CHCHD2* without affecting *Zbed5*, which has significant homology to *CHCHD2* in rodents (*24*). A silent cytosine-to-guanine mutation 2 nucleotides immediately upstream was also introduced for genotyping purpose.

**Figure 1:**
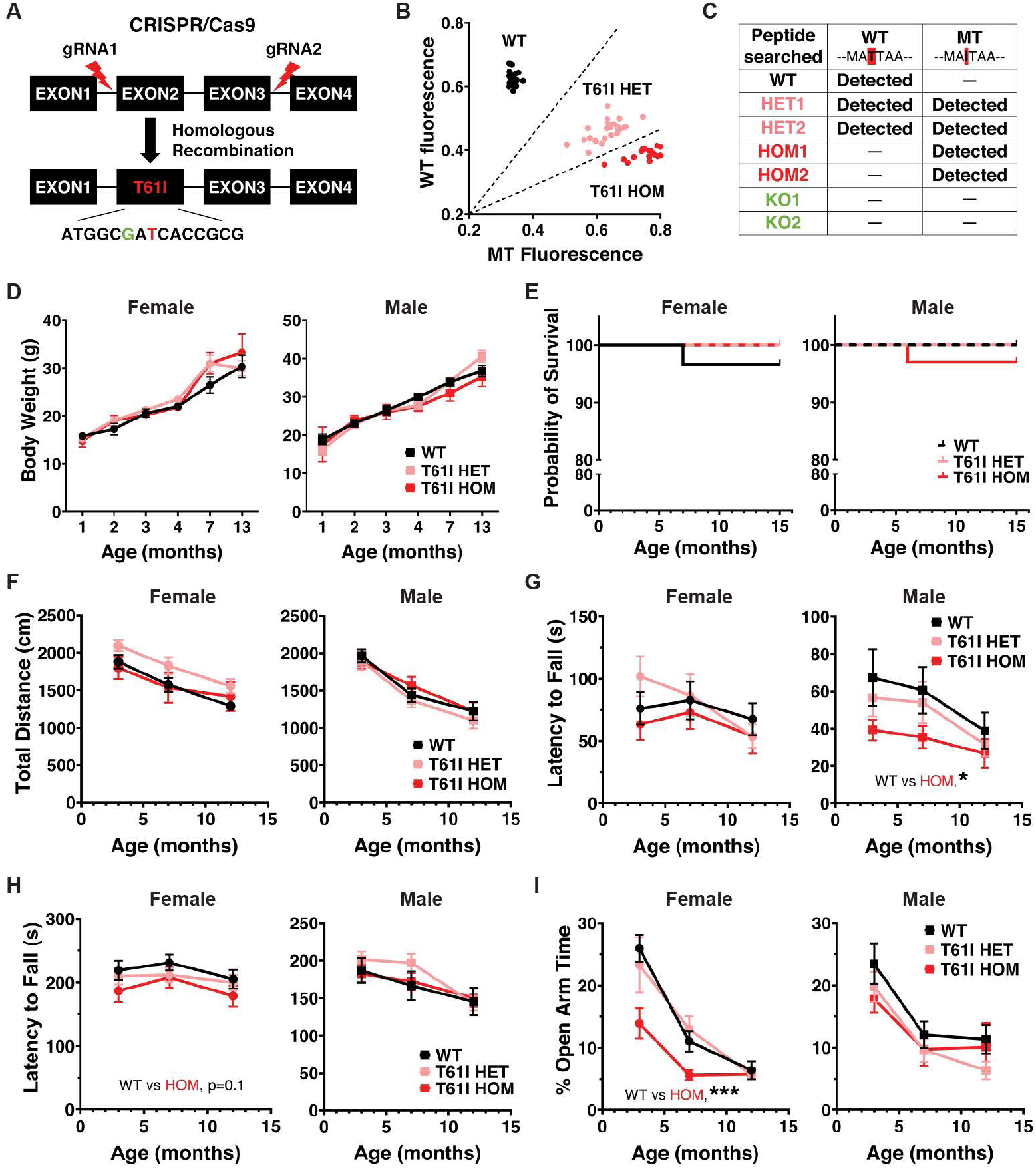
CHCHD2 T61I mice exhibit normal survival and subtle motor deficits. **(A)** Insertion of CHCHD2 T61I point mutation by CRISPR/Cas9 gene editing and homologous recombination. **(B)** CHCHD2 T61I mouse SNP genotyping by qPCR. 3 clusters were autogenerated by allelic discrimination in the analysis software, including WT (black), T61I HET (pink) and T61I HOM (red). **(C)** WT peptide (MAQMA**T**TAAGVAVGSAVGHTLGHAITGGFSGGGSAEPAKPDITYQEPQGAQL) and mutant peptide (MAQMA**I**TAAGVAVGSAVGHTLGHAITGGF) were searched in brain lysates of 7 mice of different genotypes (1 WT, 2 T61I HET, 2 T61I HOM and 2 CHCHD2 KO). WT peptide was found in WT and HET mice, while mutant was found in HET and HOM mice. Neither peptide was found in KO mice. **(D)** CHCHD2 T61I mice showed similar body weights compared with WT through 13 months of age. Data represent mean ± SEM. N = 5-9 mice/genotype/gender. **(E)** T61I mice showed normal survival compared with WT in Kaplan-Meier survival curve through 16 months of age. Left, females (n = 30 WT, 18 HET and 17 HOM); right, males (n = 21 WT, 20 HET, 34 HOM). **(F)** Total movements of T61I mice were normal in open field test. **(G)** Male T61I HOM mice showed a significant decrease in latency to fall in inverted grid hang (WT vs HOM *P* < 0.05). **(H)** Female T61I HOM mice showed a trend toward a decrease in latency to fall in the accelerating rotarod test at (WT vs HOM *P* = 0.1). **(I)** Female HOM showed significantly lower time spent in the open arms (WT vs HOM *P* < 0.001). In all behavioral studies (F-I), data represent mean ± SEM. N = 16 WT, 16 HET and 13 HOM (female), or 14 WT, 18 HET and 17 HOM (male). \**P* < 0.05, \*\*\**P* < 0.001 by linear mixed effects analysis.

Due to signal overlap from *Zbed5* in Sanger sequencing, we used qPCR to perform SNP genotyping of the mice (Fig. 1B). Different genotypes were distinguished using wild-type (WT) and mutant probes based on clustering. We also used liquid-chromatography with tandem mass spectrometry (LC-MS/MS) of total brain protein lysates as an orthogonal approach to distinguish WT and mutant peptides (Fig. 1C).

### CHCHD2 T61I mice show normal lifespan and subtle motor deficits

CHCHD2 p.T61I mice were born in normal Mendelian proportions (WT: 26.9%, HET: 42.6%, HOM: 30.6%, n=108), and had similar body weights to littermate controls through 13 months of age (Fig 1D). Survival was also unchanged through 16 months (Fig. 1E). A small subset of mice tracked for 23 months also failed to show any effects on survival (Fig. S1A). We next assessed if the CHCHD2 T61I mutation disrupts motor function. We studied both heterozygous T61I mice that models the heterozygous T61I expression observed in PD, and homozygous mutant mice which we hypothesized would have stronger phenotypes. T61I homozygous and heterozygous mutation had no overall effects on total movement in open field testing across the three time points (3, 7 or 12 months) (Fig. 1F). Repeated open field testing with the automated tracking system, EthoVision XT, on a small cohort of 23-month-old male mice also failed to show differences in total movement or time spent moving (Fig. S1, B and C). Although total movement wasn’t affected, male (but not female) homozygous T61I mice exhibited a significantly shorter latency to fall in inverted grid hang test (Fig. 1G). In rotarod testing, CHCHD2 homozygous T61I females (but not males) had a trend for decreased latency to fall versus controls (Fig. 1H). In the elevated plus maze, female T61I homozygous mutants showed a trend level decrease in distance traveled (Fig. S1D). Overall, these findings are consistent with subtle motor deficits in the T61I mice, with no evidence of progression with older age. In contrast, analysis of the motor function of CHCHD2 T61I heterozygous mice failed to uncover any significant differences across the behavioral assessments. (Fig. 1, F-H; Fig. S1, B-D).

Notably, mice with the T61I mutation also showed evidence of nonmotor changes. In elevated plus maze testing, female homozygous T61I mice spent significantly less time in the open arms, showed decreased open arm/total distance percentage, and entered the open arms fewer times, consistent with increased anxiety. (Fig. 1I; Fig. S1D).

### Electrophysiology reveals changes in dopamine autoreceptor function in CHCHD2 T61I mice

We next assessed if the T61I mutation impacts the somatodendritic physiological health of SNc DA neurons. *Ex vivo* whole cell recordings from 23-month-old male mice blind to genotype showed that basic physiological properties of SNc DA neurons in homozygous T61I mice were similar to those in control SNc neurons. As expected, most DA neurons fired spontaneously in a regular, pacemaker fashion both in cell attached and whole cell configuration (Fig. 2A-F). On average, action potentials in DA neurons from the T61I mice also had a less depolarized peak compared to controls (Fig. 2G). A parallel but not significant pattern was observed in action potential threshold (Fig. 2H). There was no difference in action potential durations (Fig. S2A). A small population of DA neurons from the T61I mice showed less regularity of firing (greater coefficient of variation of interspike intervals) compared to WT DA neurons (Fig. 2F). *I*_h_ magnitude and input resistance distributions were also similar between SNc DA neurons from WT and T61I mice (Fig. S2, B and C).

**Figure 2:**
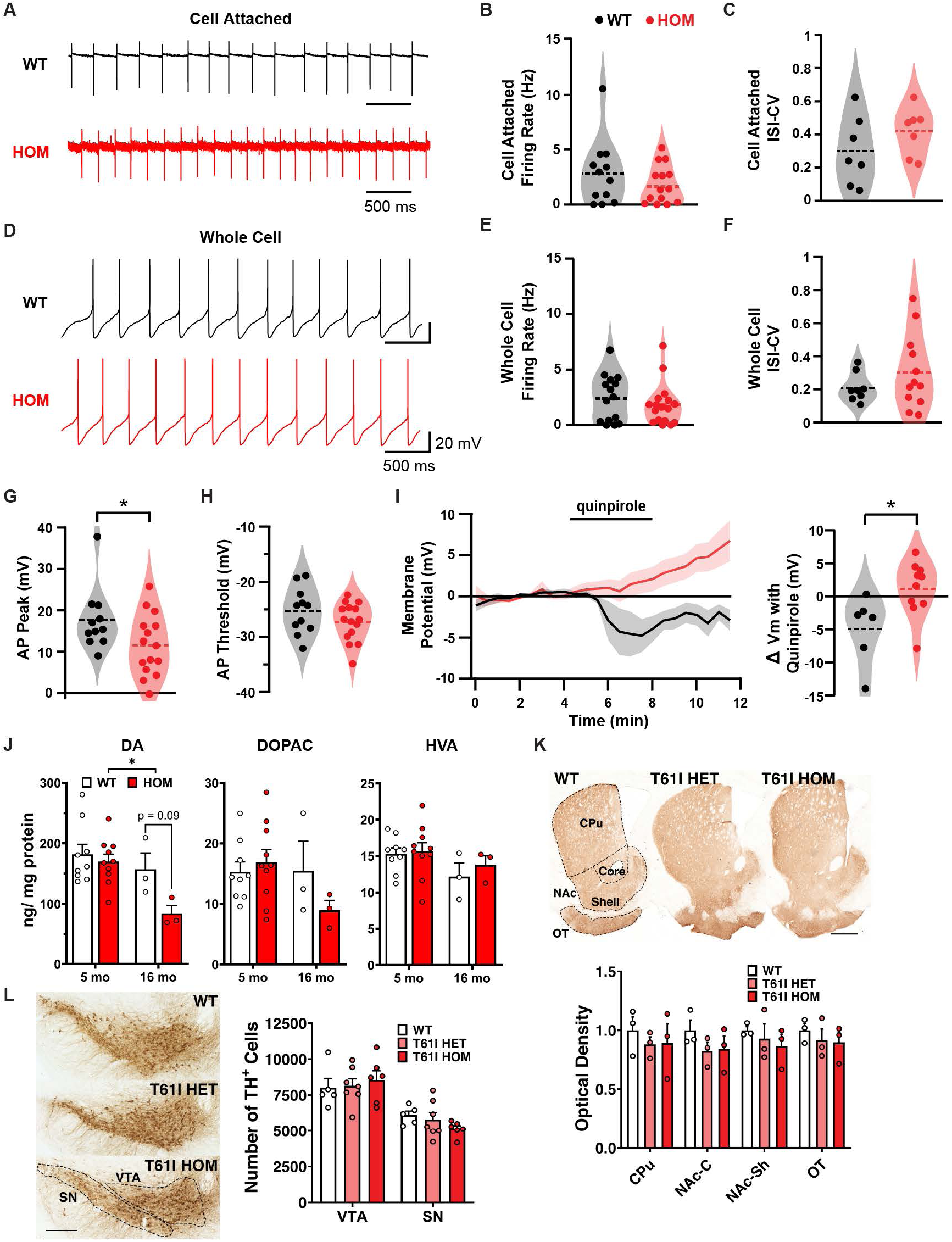
CHCHD2 T61I point mutation disrupts DA autoreceptor function without DA neuron degeneration. Recordings were made blind to genotype. **(A-C)** SNc dopamine neurons from CHCHD2 T61I and control mice fired spontaneously in acute brain slices in cell attached recording configuration. **(A)** Example action potential activity from WT and HOM mice. **(B)** Spontaneous firing rates in DA neurons from CHCHD2 T61I and control mice recorded in cell attached mode were similar. **(C)** Regularity of spontaneous firing was evaluated by measuring the coefficient of variation of 100 consecutive interspike intervals. ISI-CV was similar across genotypes. **(D)** Example pacemaker firing in DA neurons from CHCHD2 T61I and control mice in whole cell current clamp (I = 0 pA) configuration. **(E, F)** Spontaneous firing rates in whole cell current clamp configuration were similar in DA neurons from CHCHD2 T61I and control mice. **(G, H)** Spontaneous AP waveforms were recorded in whole cell configuration and quantified from each genotype including AP peak (G) and threshold (H). **(I)** The D2R selective agonist quinpirole (1 uM) was bath applied during whole cell recordings of SNc DA neurons. Quinpirole caused hyperpolarization in most WT neurons but not T61I mutant neurons. Left, time course averages of responses; right, summary of individual responses. AP, action potential. N = 5 WT and 4 HOM male mice at 23 months; each circle is a neuron. Data represent mean ± SEM or with kernel density estimations. For all data parametric assumptions were tested to choose between t-test (parametric) or permutation (non-parametric) analysis. \**P* < 0.05. **(J)** HPLC of dorsal CPu punches from flash-frozen brain tissues at 5 or 16 months showed lower DA level in T61I HOM mice. HVA, homovanillic acid; DOPAC, 3,4-dihydroxyphenylacetic acid. Data represent mean ± SEM. N = 9-10 mice/genotype at 5 months and 3 mice/genotype at 16 months. *P < 0.05 by two-way ANOVA with Sidak *post hoc* test. **(K)** Representative images of TH immunoreactivity in striatal sections of WT, T61I HET and HOM mice at 16 months. DA neuron projection areas in dorsal and ventral striatum are indicated with dotted lines. CPu, caudate putamen; NAc, nucleus accumbens; OT, olfactory tubercule. Quantification for TH immunoreactivity showed no difference in WT, HET and HOM by two-way ANOVA with Tukey’s *post hoc* test. Data represent mean ± SEM. N = 3 animals/group, 4 sections/animal. Scale bar indicates 300 μm. **(L)** Representative images of TH immunoreactivity in midbrain sections of WT, HET, and HOM mice at 16 months. DA neuron projection areas in the midbrain are indicated with dotted lines. SN, substantia nigra; VTA, ventral tegmental area. Stereology estimating the number of DA neurons showed no difference between each group by two-way ANOVA with Tukey’s *post hoc* test. N = 5-7 animals/group. Data represent mean ± SEM. \**P* < 0.05, \*\**P* < 0.01, \*\*\**P* < 0.001, Scale bar indicates 300 μm.

There was one major neurophysiological alteration in DA neuron function in the CHCHD2 T61I mice, Hikima et al (*26*) showed that dopamine release in the SNc functions specifically in an autoinhibition manner, activating D2Rs on the neuron that released the dopamine. SNc DA neurons from CHCHD2 T61I mice lacked dopamine D2 receptor (D2R) autoreceptor hyperpolarization (Fig. 2I). The lack of D2R agonist induced hyperpolarization may be due to one or more of the following factors: downregulation of D2Rs, downregulation the of G protein receptor gated inwardly rectifying K^+^ channels responsible for the hyperpolarization response, or D2R coupling changing to an alternative signaling pathway. This loss of autoinhibition probably enhances DA neuron firing activity *in vivo* in certain behavioral conditions.

To further assess the impact of the CHCHD2 T61I mutation on dopamine metabolism, we assessed catecholamine levels in the dorsal striatum. T61I homozygous mice showed lower level of DA across 5 months and 16 months (Fig. 2J), suggesting disrupted dopamine signaling.

### CHCHD2 T61I mutation does not lead to significant DA neuron degeneration

We next examined if the T61I mutation leads to DA neuron degeneration. First, to assess the integrity of DA neuron terminals, we quantified tyrosine hydroxylase (TH) expression in the striatum using 3’3-diaminobenzidine (DAB) staining. There was no significant effect of either heterozygous or homozygous T61I mutation on TH optical density in the striatum of 16 months old mice (Fig. 2K). A second cohort of 23-month-old male mice also showed no difference between the T61I and WT mice (Fig. S2D). We then performed stereology to quantify the number of DA neurons in the midbrain, and similarly there was no significant loss of DA neurons in either the SN or ventral tegmental area (VTA) (Fig. 2L).

### T61I point mutation preferentially increases the mitochondrial accumulation of CHCHD2 and CHCHD10 in SN (but not VTA) DA neurons

CHCHD2 forms soluble homodimers and heterodimers with CHCHD10 in mitochondria under physiological conditions (*27*). Moreover, homodimers and heterodimers containing T61I CHCHD2 are less soluble and more prone to aggregate (*18, 20, 28, 29*). To assess the effects of the T61I mutation on the level and spatial organization of CHCHD2, we used super-resolution microscopy on 16-month-old mice. First, we found that T61I point mutation did not change the overall level of CHCHD2 expression in either SN or VTA DA neurons (Fig. 3, A and B). Similarly, the total level of CHCHD10 expression was comparable in T61I CHCHD2 and WT controls.

**Figure 3:**
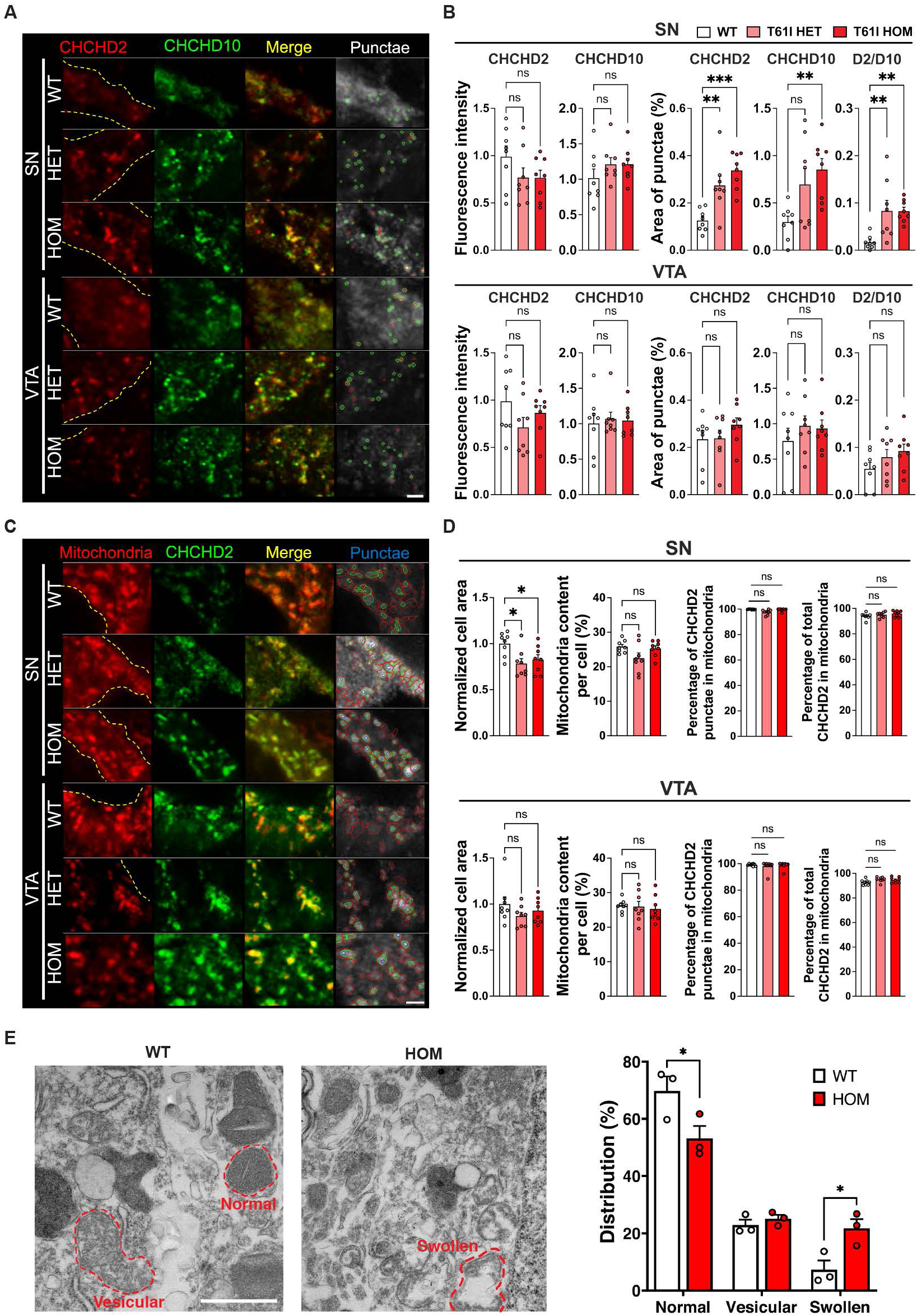
T61I point mutation preferentially increases accumulation of CHCHD2 and CHCHD10 in mitochondria in SN DA neurons. **(A)** Representative fluorescence images of CHCHD2 (red), CHCHD10 (green), and merge in SN (top) and VTA (bottom) subregions in the midbrain of CHCHD2 T61I mice at 16 months. Punctae was semi-automatically annotated by Cellprofiler. CHCHD2 denoted in red and CHCHD10 in green. The region of cell body is indicated with dotted lines, based on TH signal. **(B)** Quantification of CHCHD2 and CHCHD10 intensity, area of CHCHD2 and D10 punctae, and area of co-localization between CHCHCHD2 and D10 punctae in SN (top panel) and VTA (bottom), normalized to the averaged cell area of each group. **(C)** Representative fluorescence images of mitochondria (PDH, red), CHCHD2 (green), and merge in SN (top) and VTA (bottom). PDH denoted in red, total CHCHD2 in green and CHCHD2 punctae in blue. The region of cell body is indicated with dotted lines, based on TH signal. **(D)** Quantification of cell area, mitochondrial content, CHCHD2 punctae area in the mitochondrial, and total CHCHD2 level in the mitochondria in SN (top panel) and VTA (bottom), normalized to averaged cell area of each group. In all fluorescence staining (A-D), n = 6-8 randomly selected TH-positive cells from 6 fields in each region in each mouse, 4 mice/genotype. Examiner was blinded to mouse genotype. Data represent mean ± SEM. \**P* < 0.05, \*\**P* < 0.01, \*\*\**P* < 0.001 by one-way ANOVA with Tukey’s *post hoc* test, Scale bars indicate 1μm. **(E)** Impact of T61I mutation on mitochondrial morphology. DA neurons were labeled with primary TH antibody followed by secondary gold particle-conjugated antibody. N = 71 – 189 mitochondria/mouse, 3 mice/genotype. Data represent mean ± SEM. \*\**P* < 0.01 by *t* test with Welch’s correction. Images taken at 19,000X magnification, and scale bar indicates 1 μm.

However, both heterozygous and homozygous T61ICHCHD2-expressing DA neurons had markedly increased numbers of CHCHD2 and CHCHD10 punctae, specifically in the SN, and not the VTA. Here punctae are defined as foci of CHCHD2 or CHCHD10 immunofluorescence that are significantly brighter than adjacent areas, and hence could represent either CHCHD2 and CHCHD10 aggregation, or focal areas of increased CHCHD2 and D10 levels in mitochondria.

Since both CHCHD2 and CHCHD10 are predominantly localized to the mitochondrial intermembrane space (*28*), we next assessed if T61ICHCHD2 impacts the mitochondrial content in either SNc or VTA DA neurons. However, T61CHCHD2 heterozygous and homozygous DA neurons had similar mitochondrial content to controls, based on immunofluorescence against the inner mitochondrial marker PDH (Fig. 3, C and D). In addition, essentially all T61ICHCHD2 and wt CHCHD2 co-localized with mitochondria (Fig. 3D).

To gain insight into whether the accumulation of CHCHD2 punctae reflects a change in CHCHD2 protein solubility, for instance due to CHCHD2 aggregation, we performed western blots on cytosolic and mitochondrial fractions from total brain lysates of 16-month-old mice (Fig. S3A). T61ICHCHD2 mice had relatively increased levels of insoluble versus soluble CHCHD2 in both cytosolic and mitochondrial fractions, with the latter being more prominent, suggesting aggregation of T61ICHCHD2 within mitochondria.

### Mutant T61I CHCHD2 Disrupts Mitochondria in DA Neurons

The accumulation of T61I CHCHD2 mutant and CHCHD10 within mitochondria suggests potential disruption of mitochondria. To address whether mitochondrial cristae was affected by protein accumulation, we labeled DA neurons using immunogold staining and assessed the mitochondrial ultrastructure. Following the definition by Paredes and colleagues (*30*), we categorized mitochondrial morphology into 3 groups: normal, vesicular, and swollen. Compared with WT mice, T61I HOM mice showed a significant decrease in normal form and a significant increase in swollen form of mitochondria in DA neurons (Fig. 3E), indicating mitochondrial damage by CHCHD2 T61I mutation.

### Altered expression level of CHCHD2 in vulnerable DA neurons in PD

The change in the distribution and solubility of CHCHD2 in SNc DA neurons from T61ICHCHD2 mice suggests that changes in CHCHD2 level or conformation could contribute to disease pathophysiology. To determine whether CHCHD2 expression also changes in sporadic PD, we used spatial genomics (GeoMx WTA) to compare CHCHD2 gene expression in DA neurons in midbrain samples from patients with sporadic PD and age-matched human controls (Fig. 4A, Table S1). Interestingly, in controls, transcriptomic levels of both CHCHD2 and CHCHD10 were increased in more vulnerable DA neurons in the ventral tier of the SN (SNV), when compared with the more resilient DA neurons in the dorsal tier (SND). Considering that DA neurons in both mice and humans have increased levels of both CHCHD2 and CHCHD10 (*21*) these findings could suggest either that increased CHCHD2 contributes to the preferential vulnerability of SNV DA neurons, or that CHCHD2 is protective.

**Figure 4:**
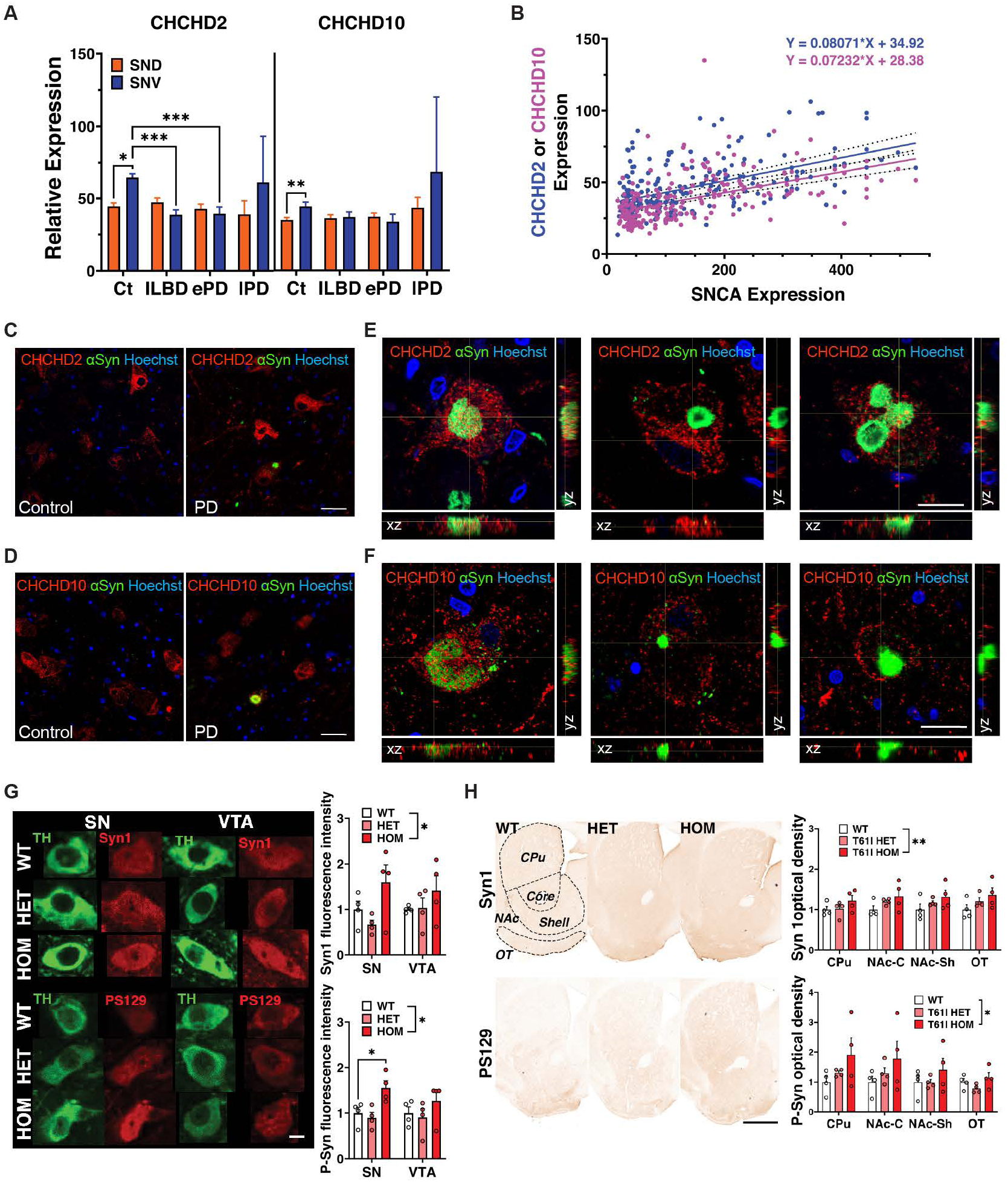
CHCHD2 T61I mice show increases in α-synuclein and phosphorylated synuclein level in the midbrain. **(A)** Transcriptomic levels of CHCHD2 and CHCHD10 in DA neurons of SND (dorsal tier of substantia nigra, orange) and SNV (ventral, blue) measured with GeoMx WTA show significant downregulation of CHCHD2 in the SNV of ILB and ePD cases. Data represent mean ± SEM. Brown-Forsythe and Welch ANOVA tests were applied for data covariates with age, sex, and post-mortem delay. \*\*\**P* < 0.001. NC, normalized counts; Ct, control; ILBD, incidental Lewy body disease; ePD, PD with early Braak stage pathology; lPD, PD with late Braak stage pathology. **(B)** Moderate positive linear associations were revealed between expression levels of either CHCHD2 or CHCHD10 with SNCA in the SN by Spearman correlation. **(C)** CHCHD2 localization in control and PD SN DA neurons and **(E)** orthogonal views of the relative location of CHCHD2 in different staged αSyn aggregations. **(D)** CHCHD10 localization in control and PD SN DA neurons, and **(F)** orthogonal views of the relative location of CHCHD10 in different staged αSyn aggregations. Scale bars represent 50 μm in (C, D), and 20 μm in (E, F). **(G)** Representative images and quantification of TH (green) and either total α-synuclein (Syn1) or phosphorylated synuclein, P-syn (PS129, red) immunoreactivity in SN and VTA at 63X magnification in midbrain sections of WT, HET and HOM mice at 16 months. Syn1 and P-syn immunoreactivities increased in midbrain DA neurons of HOM mice by two-way ANOVA with Tukey’s *post hoc* test. Data represent mean ± SEM. \**P* < 0.05. N = 4 sections/mouse, 4 mice/genotype. Scale bar indicates 10 μm. **(H)** Representative images and quantifications of Syn1 (top) and P-syn (PS129, bottom), immunoreactivity in striatal sections of mice at 16 months. Syn1 and P-Syn areas in dorsal and ventral striatum are indicated with dotted lines. Significant increases in both Syn1 and p-Syn immunoreactivity were observed in HOM mice compared with WT by two-way ANOVA with Tukey’s *post hoc* test. Data represent mean ± SEM. \**P* < 0.05, \*\**P* < 0.01. N = 4 sections/mouse, 4 mice/genotype. Scale bar indicates 300 μm.

Interestingly, in patients with early PD (ePD) there was a dramatic decrease in CHCHD2 gene expression in DA neurons in the SNV but not SND (Fig. 4A), raising the possibility that these higher CHCHD2-expressing DA neurons were lost or that CHCHD2-expression in these DA neurons decreases as they begin to degenerate. The latter possibility is supported by the finding that CHCHD2 expression is also decreased in SNV DA neurons from individuals with incidental Lewy bodies (ILB) and no significant loss of DA SNV neurons. Notably, this relationship disappeared in the late Lewy pathological stage of PD, where the resilient DA neurons remaining in the SNV tended to have higher levels of CHCHD2 expression, although the results were highly variable (coefficient of variation = 167.6%). The decrease in CHCHD2 level could reflect a downregulation of CHCHD2 expression early in PD, and the persistence of primarily higher CHCHD2-expressing DA neurons in late PD might indicate that those DA neurons with lower CHCHD2 ultimately die.

The finding of extensive α-synuclein aggregation in a patient with CHCHD2 T61I mutation suggests that mutant CHCHD2 may promote α-synuclein aggregation (*17*). To gain insight into the relationship between mutant T61I CHCHD2 and synuclein, we first examined the relationship between CHCHD2 and *SNCA* gene expression by spatial transcriptomics. As with CHCHD2, *SNCA* expression in controls was higher in DA neurons in the most PD-vulnerable ventral tier (SNV, 238.80 ± 131.32) than those in the dorsal tier (SND, 130.78 ± 130.86, *P* < 0.001). Considering the known pathogenicity of increased WT α-synuclein (*31*), this suggests that increased α-synuclein levels may contribute to the preferential vulnerability of ventral tier DA neurons. Interestingly, when pooling all samples (Fig. 4B), regression tests indicated positive linear associations between the level of SNCA and CHCHD2 (R2 = 0.270, F (1,226) = 83.74, *P* < 0.001) as well as between the level of SNCA and CHCHD10 (R2 = 0.279, F (1,226) = 87.51, *P* < 0.001).

### CHCHD2 and CHCHD10 accumulate in **α**Syn aggregates in SNc dopamine neurons in sporadic PD

To further examine the relationship between CHCHD2 and α-synuclein, we next compared CHCHD2 localization in SNc DA neurons from PD and age-matched controls. As observed in mouse brains, both CHCHD2 and its binding partner CHCHD10 had a predominantly punctate pattern within the cytoplasm of SNc DA neurons in both control and PD patients (Fig. 4, C and D), indicating their mitochondrial localization. In PD samples, there was a robust accumulation of CHCHD2 and CHCHD10 with α-synuclein (assessed in 74 Lewy aggregates for CHCHD2 and 88 Lewy aggregates for CHCHD10). CHCHD2 and CHCHD10 immunoreactivity was always either embedded in the Lewy aggregates or at the edge of mature Lewy bodies (Fig. 4, E and F).

Given the above correlations between CHCHD2 and α-synuclein gene expression in SNc DA neurons and the distribution of CHCHD2 and CHCHD10 in α-synuclein aggregates in PD cases, we next examined if α-synuclein accumulates and aggregates in the CHCHD2 T61I mutant mice. We first performed western blots on the cortex of 16-month-old mice. However, there was no change in α-synuclein levels in either the serially extracted Triton-soluble or SDS-soluble fractions (Fig. S3B).

To assess if the T61I mutation impacts synuclein levels specifically in midbrain DA neurons, we next assessed α-synuclein and phosphorylated synuclein (P-Syn) levels specifically in midbrain DA neurons using immunofluorescence (Fig. 4G). Quantification revealed an increase in P-Syn levels in homozygous SNc DA neurons. Non-phosphorylated α-synuclein levels were also increased overall in homozygote midbrain DA neurons. Moreover, both α-synuclein and phosphorylated synuclein were increased in the striatum of homozygous T61I mice at 16 months by optical density, while heterozygotes did not reach significance (Fig. 4H). Therefore, T61I homozygous mice accumulate increased levels of both total and phosphorylated α-synuclein in both the cell body and striatum. However, further investigation is required to determine if SNc DA neurons in heterozygous T61I mice also accumulate increased α-synuclein at older age, and also whether the increase in the striatum reflects changes in midbrain DA axons versus striatal target neurons. Similarly, it will be important to clarify whether the lack of accumulation in cortical brain lysates truly reflects a selective predisposition to accumulate α-synuclein in midbrain DA neurons and the striatum, or simply reflects a lower sensitivity of the western blot approach to detect changes in α-synuclein level. Further studies are needed to understand the mechanisms by which mutant CHCHD2 affects synuclein levels in midbrain DA neurons.

### CHCHD2 mutant mouse brains exhibit a metabolic shift toward glycolysis and the pentose phosphate pathway (PPP)

The accumulation of T61I CHCHD2 mutant and CHCHD10 within mitochondria, as well as the disruption of mitochondrial morphology suggests that mutant T61I CHCHD2 mutant may disrupt mitochondrial function. Previous studies have highlighted a metabolic switch from mitochondrial respiration to glycolysis as a compensatory mechanism to sustain cellular ATP in the context of mitochondrial dysfunction (*32*). Notably, recent work with a DA neuron-specific complex I deficiency mouse model revealed a robust upregulation of glycolytic genes in DA neurons by RNAseq (*33*). Similarly, pronounced elevation of both glycolytic gene mRNA and protein levels were also found in the heart of CHCHD10 point mutant mice (*34*). To determine if mutant CHCHD2 disrupts energy metabolism, we performed targeted and untargeted metabolomics where mice were infused with [U-^13^C]glucose for 30 minutes before harvest.

Remarkably, female T61I heterozygous and homozygous mice at 19 months exhibited a robust dose-dependent increase in the total level of distal glycolytic metabolites including 3-phosphoglycerate (3PG), phosphoenolpyruvate (PEP) and pyruvate (Pyr) (Fig. 5, A and B). The fractional labeling with 13C was also increased across essentially all glycolytic metabolites in homozygous mice versus controls, consistent with an overall increase in brain metabolism of glucose through glycolysis (Fig. 5, A and C). T61I heterozygous and homozygous mice also showed dose-dependent increases in the fractional labeling of PPP metabolites and in relative amounts of a subset of TCA cycle metabolites. However, fractional labeling of TCA metabolites was only subtly increased in homozygous mice versus controls, suggesting that the accumulation of TCA metabolites occurs due to a decrease (rather than increase) in the flux of glucose metabolites toward respiration. This overall increase in glucose metabolism through glycolysis and the PPP relative to the TCA cycle is thus likely secondary to a primary disruption in mitochondrial respiration.

**Figure 5:**
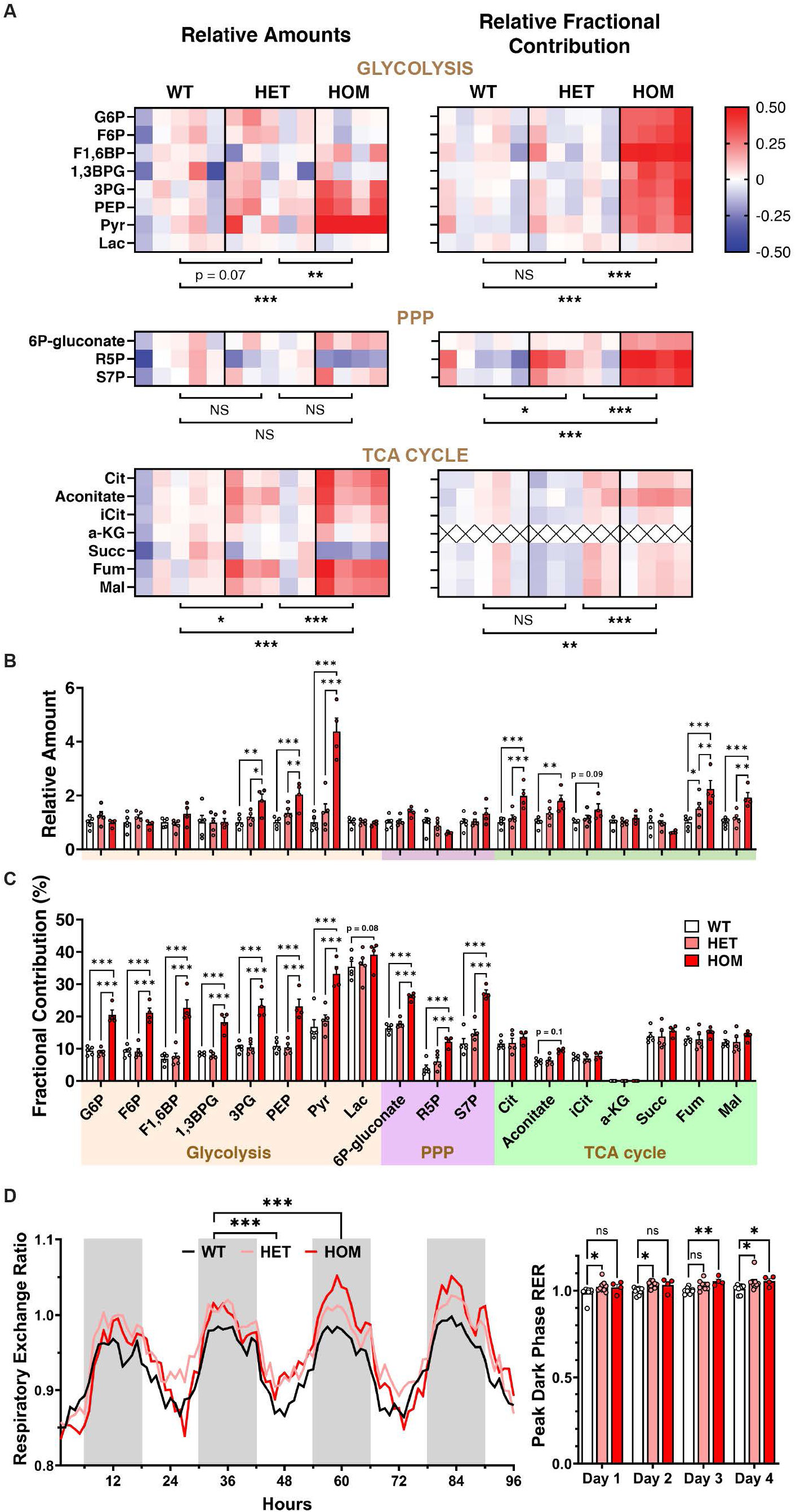
CHCHD2 mutant mouse brains exhibit a metabolic shift toward glycolysis and PPP. **(A)** Female CHCHD2 T61I mice at 19 months received tail vein injections of [U-^13^C]glucose for 30 minutes before brains were harvested. Brain metabolite extract was analyzed by an ion chromatography-mass spectrometry (IC-MS) detector. Heat map shows relative total levels of metabolomics in the L column determined by non-targeted metabolomics, and fractional contribution of 13C-glucose to each metabolite by targeted metabolomics on the R. Each square shows the log value of a metabolite level or fractional contribution in a mouse normalized to the mean of the WT group. **(B)** Bar graph of relative amounts of metabolites along glycolysis, PPP and TCA cycle. CHCHD2 T61I heterozygotes and homozygotes showed a dose-dependent increase in distal glycolytic, PPP and TCA cycle metabolites compared with WT mice. **(C)** Bar graph of fractional contribution of metabolites. CHCHD2 T61I homozygous mice showed significantly higher labeling percentages in glycolytic and PPP metabolites compared with WT mice. G6P, glucose 6-phosphate; F6P, fructose 6-phosphate; F1,6BP, fructose 1,6-bisphosphate; 1,3BPG, 1,3-bisphosphoglycerate; 3PG, 3-phosphoglycerate; PEP, phosphoenolpyruvate; Pyr, pyruvate; Lac, lactate; R5P, ribose 5-phosphate; S7P, sedoheptulose 7-phosphate; Cit, citrate; iCit, isocitrate; a-KG, α-ketoglutarate; Succ, succinate; Fum, fumarate; Mal, malate. Data represent mean ± SEM. N = 5 WT, 5 HET and 4 HOM T61I female mice at 19 months. \**P* < 0.05, \*\**P* < 0.01, \*\*\**P* < 0.001 by two-way ANOVA with Tukey’s *post hoc* test. **(D)** Respiratory exchange ratio (CO_2_ production to O_2_ consumption) of CHCHD2 T61I mice at 5 months was recorded for 5 days. T61I HOM mice showed significant increase in RER in the dark phase. N = 9 WT, 9 HET and 4 HOM. Data represents mean ± SEM in the bar graph. \**P* < 0.05 by two-way ANOVA with Dunnett’s *post hoc* test.

### CHCHD2 T61I mice show increased carbohydrate consumption

Given our finding of the metabolic shift toward glycolysis, whole-body metabolism of the T61I mice at 5 months was monitored by the comprehensive lab animal monitoring system (CLAMS). Throughout the 5 days in CLAMS, both heterozygous and homogygousT61I mice showed an increased respiratory exchange ratio (RER), particularly in the dark phase when mice were more active (Fig. 5D). The decrease in O_2_ consumption per carbon implies a fuel shift toward carbohydrate consumption for glycolysis without affecting overall energy expenditure (Fig. S3C), in line with the disruption in respiration implied by metabolomics. As the measurements of whole-body metabolism primarily reflect changes from tissues outside of the brain, especially skeletal muscle and adipose tissues, this data shows that CHCHD2 T61I mice have systemic metabolic changes that extend beyond the brain.

### Spatial transcriptomics reveals down-regulated respiratory chain gene expression in DA neurons in T61I mice

To investigate how CHCHD2 impacts gene expression in DA neurons, we performed spatial genomics (Visium Spatial Gene Expression,10X Genomics) on midbrain sections from 16-month-old CHCHD2 T61I and from 13-month-old CHCHD2 KO mice. Disks demarcating areas of gene expression analysis were selected within the SN and VTA, based on containing at least one entire TH-positive neuron and also expressing characteristic DA genes (DAT, VMAT2, or TH) above a pre-set threshold level (Fig. S4, A and B). We first examined selected pathways, including glycolysis, pentose phosphate pathway, respiratory chain and integrated stress response. Both T61I CHCHD2 and knockout showed significant effects on the metabolic pathways (Fig. S4, C and D). The glycolytic pathway was downregulated in the VTA of CHCHD2 KO mice, while unaffected in CHCHD2 T61I mice (Fig. S4E). In contrast, expression level of nearly every component in complex I and III was decreased in the VTA of CHCHD2 T61I mice, but had subtle effects in the KO mice (Fig. S4, F and G).

We then performed untargeted analyses to identify individual hits and pathways in these mice. Genes ranking within the top 20 percent of significantly altered ones in either the SN or VTA of the CHCHD2 T61I mice were carefully reviewed (Fig. S5A). Surprisingly, none of these genes was within the top 20 percent hits in the SN and VTA of CHCHD2 KO mice, and vice versa (Fig. S5B). This implies T61I CHCHD2 exerts different mechanisms than that of CHCHD2 KO. Among genes with the most pronounced changes in CHCHD2 T61I mice (Fig. S5A), we identified three shared hits in both the SN and VTA: *Zfand6*, *Lrmda* and *Apobec1* (Fig. S5C). Zfand6 (zinc finger AN1-type containing 6, also known as AWP1), is a key mediator of TNF-α signaling and negatively regulates NF-κB (*35–37*). Little is known about Lrmda (leucine rich melanocyte differentiation associated, also known as C10orf11), but mutations in this gene cause oculocutaneous albinism. Apobec1 is a cytidine deaminase enzyme which edits transcripts in various tissues including macrophages and dendritic cells. Deletion of Apobec1 in microglia in mice can exacerbate age-related neurodegeneration (*38*). These genes did not show similar trends in the CHCHD2 KO mice versus their littermate controls. Additionally, we also performed pathway analysis on the CHCHD2 T61I mice using the GSEA tool, but none passed the false discovery threshold of 0.25.

### Human spatial transcriptomics reproduces similar changes in selected genes in mouse SN DA neurons

To assess if gene expression changes in the SN and VTA of CHCHD2 T61I mice are replicated in the SNc in sporadic PD, we examined the expression levels of DEGs (differentially expressed genes) in ILB cases versus healthy controls (Fig S6). Since the demarcating disks included both DA neurons and surrounding cells in the mouse brains due to the limited resolution of Visium, we performed whole tissue analysis in the human samples without TH+ masking. Out of the 17 DEGs, only 8 were within our GeoMx human dataset assessing gene expression within SNV TH+ cells. Remarkedly, 7 of these genes exhibited changes in the same direction as observed in the mouse model, and one (Cox6a2) reached significance. This data supports the conclusion that CHCHD2 T61I point mutation leads to transcriptomic changes similar to those in sporadic PD.

### Proteomics reveals robustly altered mitochondrial protein-protein interactions in the CHCHD2 T61I mice

To explore how mutations in CHCHD2 might impair energy metabolism and contribute to the aggregation of CHCHD2, we investigated the protein-protein interactions (PPI) of total proteasome in the brains of 13-month-old CHCHD2 T61I HET, T61I HOM, and CHCHD2 KO mice. We utilized co-fractionation coupled with mass spectrometry (CF-MS) to identify changes in protein complexes within whole cell and mitochondrial extracts from CHCHD2 point mutant and knockout (KO) mice brains (Fig. 6A). We analyzed a total of 768 fractions using size-exclusion high-performance liquid chromatography (SEC-HPLC) followed by tandem liquid chromatography-mass spectrometry/mass spectrometry (LC-MS/MS). Following the fractionation experiments, we observed the molecular weights deduced from SEC chromatographic elution profiles (Fig. S7A) were consistent. Using hierarchical clustering on the elution profiles from both whole cell lysates (WCL) and mitochondrial extracts, as determined by MS/MS, we successfully identified 9,300 proteins from the mouse proteome in WCL and 297 out of 1,140 mitochondrial proteins listed in the mouse MitoCarta database in the mitochondrial extracts (Fig. 6B). This analysis revealed distinct patterns of protein assemblies that consistently co-eluted in specific extracts. Notably, proteins involved in oxidative phosphorylation and linked to Parkinson’s disease were found exclusively in mitochondrial extracts, but not in whole cell lysates (Fig. S7B).

**Fig. 6.**
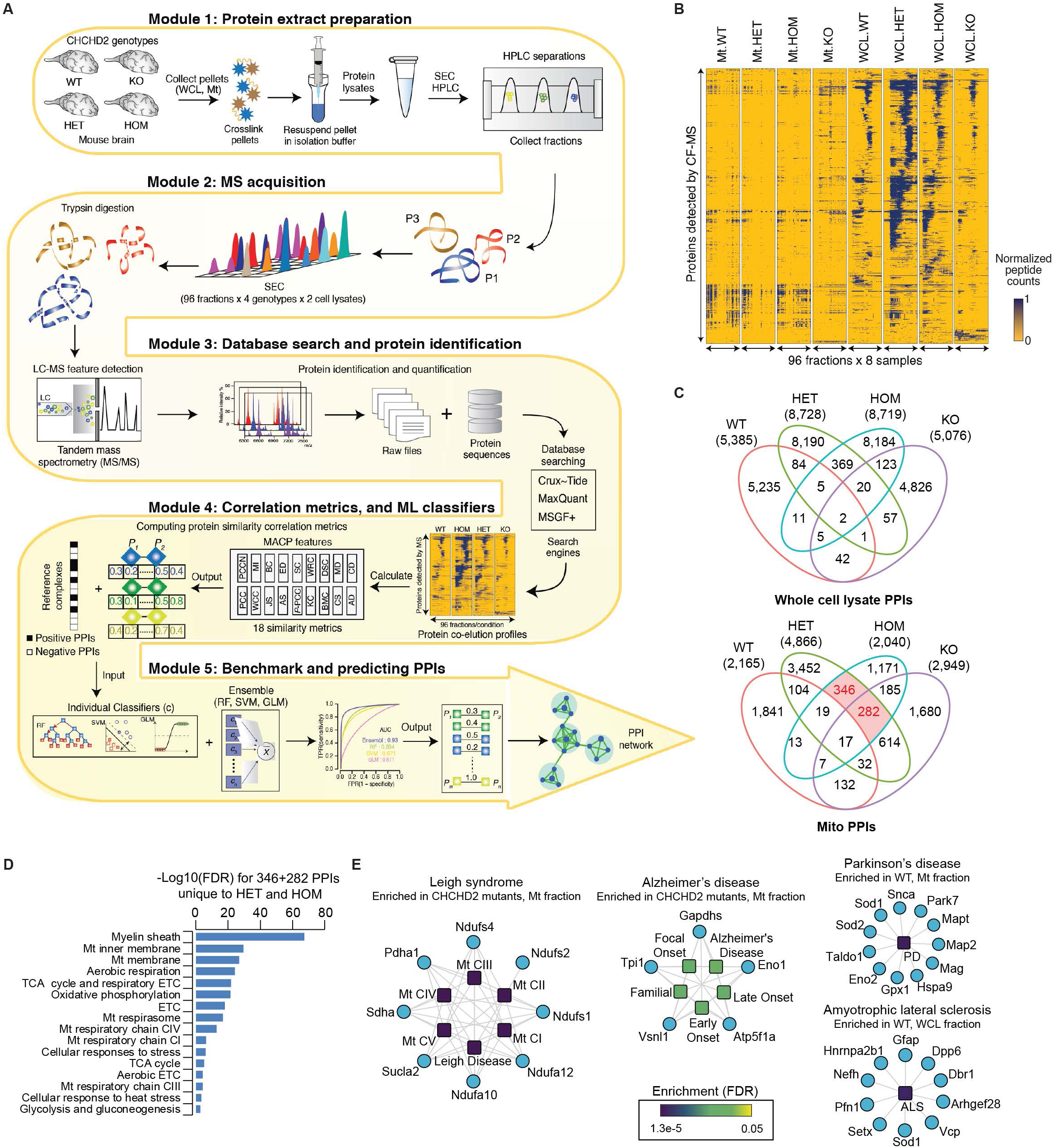
Co-fractionation mass spectrometry analysis of protein assemblies across various CHCHD2 genotypes. **(A)** Biochemical fractionation of whole cell lysates (WCL) and mitochondrial (mt) lysates from mouse brain with varying CHCHD2 genotypes was performed using high-performance size exclusion chromatography (SEC-HPLC). Protein fractions were subjected to tryptic digestion and analyzed by high performance liquid chromatography-tandem mass spectrometry (LC-MS/MS) to measure peptide spectral counts. The schematic details the computational scoring pipeline from our previous work in MACP. This pipeline includes calculating protein similarity (correlation) metrics for each CHCHD2 genotype, training integrative classifiers with machine learning using the CORUM mouse complex database as a training standard, and scoring co-fractionation data to predict high-confidence interactions. These predicted interactions were clustered to define co-complex membership, and analyzed for pathobiological relevance. **(B)** Hierarchical clustering displaying changes in protein co-fractionation intensity profiles between mt and WCL as measured by LC-MS/MS. **(C)** Venn diagram illustrating the distribution of mouse protein interactions within the mt and WCL across different CHCHD2 genotypes. **(D)** Enrichment analysis of interacting proteins in mt mouse brain lysates from CHCHD2 heterozygous (HET) and homozygous (HOM) samples involved in annotated metabolic processes. **(E)** Enrichment analysis (FDR ≤ 5e-02) of protein assemblies in mt extracts from CHCHD2 point mutants, not found in WT, and also present in both whole cell and mitochondrial extracts of CHCHD2 WT but missing in mutants. These assemblies are linked to diseases such as Leigh syndrome, Parkinson’s disease, amyotrophic lateral sclerosis, and Alzheimer’s disease.

Utilizing our newly developed R/CRAN software (*39*), Macromolecular Assemblies from Co-elution Profiles (MACP), we calculated correlation scores for each co-fractionation experiment. We generated MS2 spectral count profiles across all fractions for each genotype within each cell extract, employing three distinct search engines (Tide, MSGF+, MaxQuant). These were then analyzed using 18 different elution profile similarity metrics (Fig. 6A). By aggregating the biochemical correlation scores from these experiments, we assessed the accuracy of various machine learning classifiers—individual (random forest generalized linear model, support vector machine) and ensemble— in predicting protein-protein interactions (PPIs) through five-fold cross-validation, referencing CORUM mouse protein complexes as our benchmark. The ensemble classifier demonstrated superior sensitivity over individual models, identifying a greater number of CORUM protein pairs without compromising specificity (Fig. S7C). Setting a PPI score threshold between 0.79-0.94 for whole cell lysate and 0.75-0.84 for mitochondrial fractions based on the auROC of the ensemble model, and after refining the dataset by filtering out unlikely connections (i.e., between outer mitochondrial membrane and matrix as well as between intermembrane space and matrix), MACP identified a robust set of 5,076-8,728 PPIs from whole cell lysate and 2,040-4,866 PPIs from mitochondrial fractions, involving 668-1,356 and 455-818 proteins, respectively (Fig. 6C; Fig. S7D). Particularly, GO biological function analysis (Fig. 6D) on mitochondrial fraction from CHCHD2 mutant-exclusive intersections (highlighted in Fig. 6C) revealed significant enrichment (FDR = 5.0e-02) for metabolic processes, including mitochondrial respiration and glycolysis, among others.

Notably, protein assemblies identified in mitochondrial extracts from CHCHD2 point mutants, but not inWT, as well as those identified in both whole cell and mitochondrial extracts of CHCHD2 WT but not in mutants, showed a marked enrichment (FDR ≤ 5e-02) for associations with diseases such as Leigh syndrome, Parkinson’s disease, amyotrophic lateral sclerosis and Alzheimer’s disease (Fig. 6E).

Following the establishment of the PPI network for each genotype, the ClusterONE algorithm was employed to divide these networks into numerous distinct protein complexes within the mouse brain’s whole cell and mitochondrial lysates. The components of these predicted complexes for each genotype were then consolidated into a gene set. This gene set was compared against the co-elution profiles to pinpoint differential complexes that showed enrichment in the CHCHD2 mutants or in WT (Fig. 7, A and C). We identified PPIs altered in opposite directions by CHCHD2 KO or T61I mutation, particularly in the mitochondrial fraction (Fig. 7C), supporting the gain-of-toxic-function hypothesis. We then focused on identifying common proteins across complexes in each cluster, and observed decreased interactions with glycolytic enzymes (cluster 1) and increased interactions with ribosomal (cluster 2) and proteasomal (cluster 3) proteins in whole-cell lysates of both CHCHD2 T61I and KO mice (Fig. 7B). Notably, a significant increase in respiratory chain component interactions was found in the mitochondrial fraction in both T61I and KO mice (Fig. 7D), suggesting the crucial role of CHCHD2 in maintaining respiratory chain assembly. In examining distinct patterns in T61I HOM compared to T61I HET and KO mice (cluster 2), we highlighted Dnajc11, which plays an important role in maintaining mitochondrial cristae structure and may be associated with neurodegenerative diseases. We also observed a potential association between mutant CHCHD2 and other genes related to neurodegenerative diseases, including DJ-1 (Park7) in cluster 3, which leads to an autosomal recessive form of PD when mutated. Others include Chrnb4 in cluster 3 and Cldn17 in cluster 4 both associated with frontotemporal dementia, Nefh in cluster 5 with amyotrophic lateral sclerosis, and Vdac2 in cluster 6 with Alzheimer’s disease. These findings revealed robust disruptions in cellular metabolism due to CHCHD2 dysfunction and suggested potential linkages of mitochondrial malfunction to neurodegeneration.

**Fig. 7.**
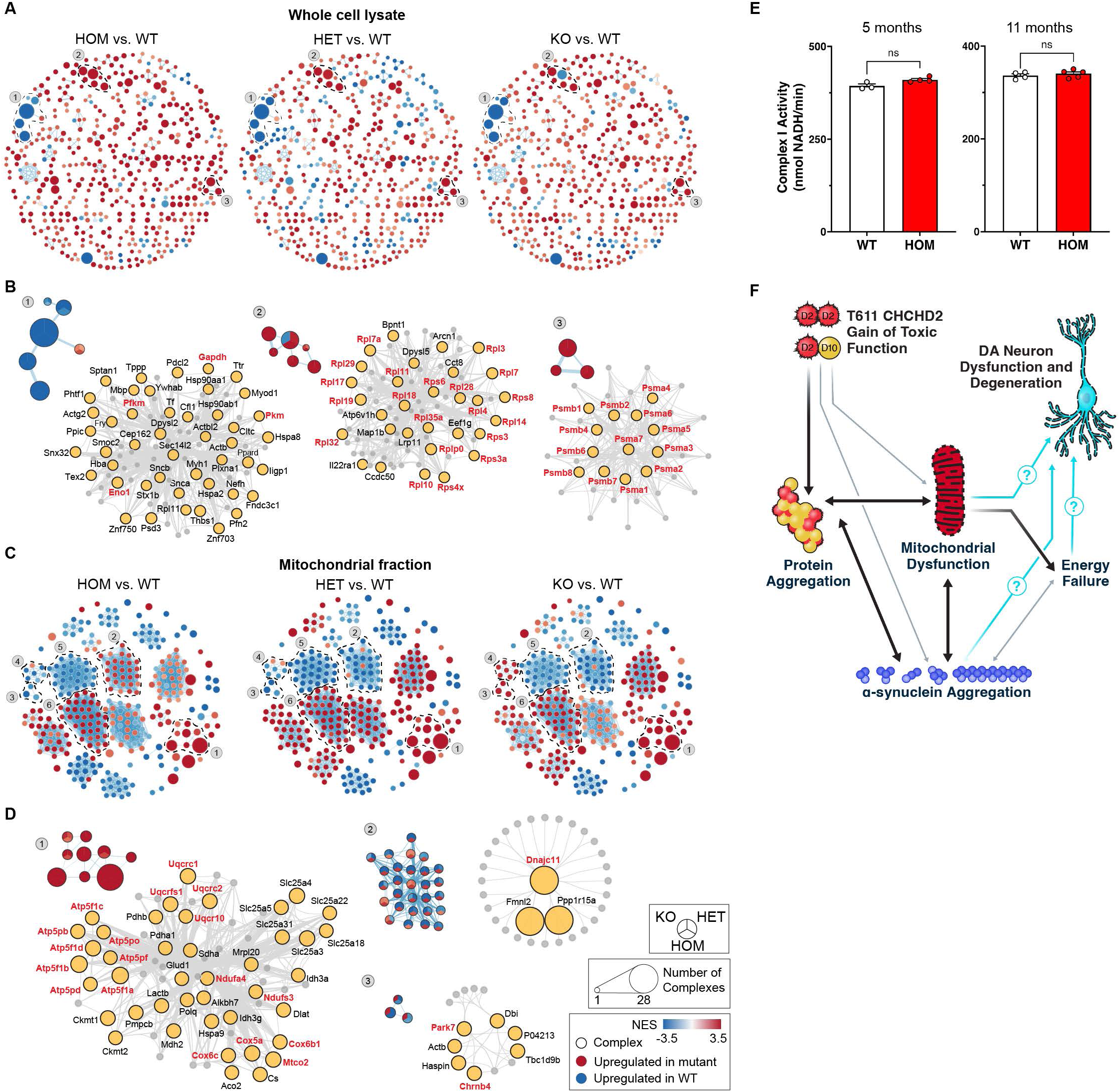
Differential analysis of predicted protein complexes in mouse brain WCL and mt lysates across different CHCHD2 genotypes. **(A, C)** Complexes in WCLs (A) and mt lysates (C) exhibit differential enrichment between CHCHD2 mutants and WT. **(B, D)** Displays of representative complexes (indicated by dotted circles with labels) showing increased interactions in WCLs (B) and mt lysates (D) of either CHCHD2 mutants or WT. The left panels in B and D show the proportion of interactions within each node, detected across various CHCHD2 genotypes or WT, with node size representing the number of interacting proteins predicted in each complex. The right panels in B and D provide detailed views of the physically associated proteins within each cluster. **(E)** Complex I activity was not changed in T61I HOM mice at 5 and 11 months. N = 3 WT and 4 HOM mice at 5 months and 4 WT and 5 HOM mice at 11 months. Data represents mean ± SEM. ns, not significant by t test with Welch’s correction. **(F)** Summary schema of the study.

### Complex I of homozygous CHCHD2 T61I mice shows normal activity

As both spatial transcriptomics and proteomics data suggest a disruption in respiratory chain complex I components, we tested complex I activity by measuring NADH consumption in 45 minutes (Fig. 7E). Isolated mitochondria from female T61I homozygous mice at 5 months old showed trending higher activity compared with WT mice, but this difference was concealed at 11 months. This suggests the respiratory disruption may not result from blockages of a single step, but an overall down-regulated performance of the whole pathway.

## DISCUSSION

Our understanding about how mitochondrial dysfunction can cause PD is limited by the lack of robust phenotype in mitochondria-based monogenic mouse models of PD including Parkin KO, PINK1 KO and DJ-1 KO (*40*). This may not be surprising given the short mean lifespan of mice, while Parkin, PINK1 and DJ-1 based PD usually begins after age 20 (*41*). However, both Parkin and PINK1 PD have been reported in patients 5 or younger (*42, 43*), suggesting that other fundamental differences between mice and humans may also contribute to the lack of robust phenotype.

In contrast, even though most patients with T61I CHCHD2-associated PD to date haven’t presented until their 50s (*16*), and idiopathic PD typically starts in the 60s, T61I CHCHD2 mice develop profound CHCHD2 aggregation, resembling the aggregation reported in a CHCHD2 PD patient (*17*). Interestingly, these CHCHD2 aggregates were reported to have decreased colocalization with a mitochondrial marker (ATP5A), and hence interpreted as accumulating in the cytosol, although the percent localizing to the cytosol was not quantified (*44*). In contrast, in our mouse model, we observe near complete co-localization of CHCHD2 aggregates with mitochondria, perhaps suggesting that the aggregates first start in the mitochondria, but then a fraction escape to the cytosol. Alternatively, undefined technical or species differences may underlie this discrepancy. We hypothesize that the accumulation of CHCHD2 aggregates lead to disruption of mitochondrial morphology and mitochondrial protein-protein interactions, ultimately leading to mitochondrial dysfunction and a consequent metabolic shift that may be common features by which mitochondrial dysfunction causes different subtypes of both genetic and sporadic PD. Similarly, T61I CHCHD2 mice also show an accumulation of total and phosphorylated synuclein in midbrain DA neurons, mimicking changes that occur in both CHCHD2-PD (*17*) and idiopathic PD.

### CHCHD2 T61I Mouse Models of PD

Two other similar CHCHD2 T61I knock-in mouse models have been reported recently, producing somewhat divergent results from ours. The T61I knock-in mouse reported by Shimizu et al. demonstrated subtle impairments on rotarod in a very small cohort and decreased TH fluorescence signal in the SN at 30 weeks, although did not directly assess DA neuron degeneration (*24*). In contrast, the other T61I CHCHD2 mouse reported by Fan et al. exhibited decreased body weight from 44 weeks, accompanied by a substantial decrease in mouse survival starting at 25 weeks, suggesting undetermined fundamental differences in these mice from our model. The weight loss and decreased mouse survival was accompanied by ≈30% decrease in the number of TH positive neurons in the SN but not VTA at 11 months, although stereology was not used (*25*). Notably, cell loss was not confirmed with another marker, raising the possibility that the decrease in TH neurons reflected downregulation of TH expression rather than cell death. Indeed, a substantial loss of TH neurons would be surprising given the subtle decrease in striatal TH optical density suggesting little if any loss of terminals (*25*), and would contrast with prior mitochondrial-based models targeting DA neurons where axonal loss precedes neuronal death (*33*). Notably, a third study involving transgenic overexpression of human T61I CHCHD2 on a background of WT CHCHD2 using a prion promoter (*44*) also reports substantial decrease in TH cell counts in the SN with no change in the VTA, assessed by immunofluorescence and a subtle decrease in striatal TH intensity, perhaps reflecting increased toxicity of higher levels of the mutant T61I CHCHD2, although the quantification methods are not provided, and it’s also again unclear if this reflects downregulation of TH rather than true DA neuron loss.

### Mutant CHCHD2 Disrupts Mitochondrial Respiration

CHCHD2 and its binding partner CHCHD10 have been shown to support mitochondrial respiration and mediate integrated stress response in different models (*18, 19*, *20, 45, 46*). We hypothesize that T61ICHCHD2 ultimately promotes neurodegeneration by disrupting energy metabolism. Although this remains to be proven, several lines of evidence from our mouse model support this hypothesis. First, CHCHD2 is primarily a mitochondrial cristae protein, and we show that the T61ICHCHD2 mutation promotes the accumulation of CHCHD2 in the mitochondria in DA neurons. Second, T61ICHCHD2 disrupts the morphology of mitochondria in DA neurons, suggesting that their function is likely compromised. Third, spatial genomics reveals subtle but widespread decreases in the expression of mitochondrial complex I and III genes in midbrain DA neurons. Fourth, untargeted metabolomics reveal an accumulation of both TCA and glycolytic metabolites consistent with mitochondrial dysfunction, with increased fractional labeling of glycolytic metabolites again consistent with a primary disruption of aerobic respiration. Fifth, whole mouse calorimetry studies reveal proportionally more reliance on carbohydrates relative to fat, also consistent with a deficit in aerobic respiration. Last, analysis of mitochondrial protein-protein interactions in brain lysates shows a broad disruption of interactions between respiratory chain proteins in the mouse brain.

We hypothesize the aggregation of CHCHD2 plays a causal role in the disruption of respiratory and other mitochondrial protein-protein interactions in CHCHD2 mutant mice. An alternate possibility is that a primary disruption of mitochondrial protein-protein interactions by T61ICHCHD2 leads to the disruption of energy metabolism, and the aggregation and accumulation of both T61ICHCHD2 and CHCHD10 are both secondary processes that do not influence (or even protect) against the degeneration process. Similarly, we hypothesize that the metabolic dysfunction leads directly to the DA neuron dysfunction and (in humans) neurodegeneration, but these steps also need to be established.

Our proteomic analyses reveal other potentially important interactions. Among these, multiple changes in the PD protein DJ-1 were observed, raising the possibility of a mechanistic connection between DJ1 and PD. DJ-1 was recently reported to directly bind to the CHCHD2 promoter and negatively regulate its expression (*47*). In addition, Woo and colleagues also observed a decrease in DJ-1 solubility in the midbrain of CHCHD2 T61I by western blotting(*44*). CHCHD2 and DJ-1 both can function in the integrated stress response and mitochondrial homeostasis, particularly in response to oxidative stress (*48, 49*), suggesting that they may normally act together to maintain mitochondrial quality control.

### CHCHD2 Mutation Can Cause **α**-Synuclein Aggregation

Mitochondrial dysfunction and increased α-synuclein both cause rare monogenic forms of PD, and both also occur in idiopathic PD. In addition to producing prominent changes in mitochondria and energy metabolism, our T61I CHCHD2 mouse model also produces increased levels of both total and phosphorylated-S129 α-synuclein, consistent with findings in the other two CHCHD2 KI mouse models (*24, 25, 44*). Combined with the prominent α-synuclein aggregation observed in a patient with T61I CHCHD2, this provides compelling evidence that a PD-causing mitochondrial mutation can cause α-synuclein aggregation. This is important in strengthening a causative link between mitochondrial dysfunction and typical later onset PD, especially because the lack of Lewy pathology in most patients with Parkin-based PD has led many investigators to conclude that autosomal recessive forms of PD are distinct from sporadic PD.

Our model system also provides opportunity to study selective vulnerability. We previously showed that midbrain DA neurons have elevated levels of CHCHD2 relative to several other neuron types(*21*), suggesting a potential mechanism for selective vulnerability. Here, we show that SNc DA neurons preferentially accumulate CHCHD2 aggregates versus VTA DA neurons. Interestingly, we also show that gene expression of CHCHD2 is elevated specifically in vulnerable DA neurons in the ventral tier of the SNc, relative to levels in the dorsal tier. In contrast, this difference was not detected in either pre-symptomatic cases with incidental Lewy bodies where there is no significant loss of DA neurons or in early PD, suggesting that it is unlikely to reflect the death of DA neurons with higher CHCHD2. Instead, it may reflect a downregulation of CHCHD2 levels in DA neurons as the disease progresses, which could indicate that it normally protects or is toxic. However, we also observed that CHCHD2, together with mitochondria, are engulfed in early Lewy aggregates, and presumably degraded as they are no longer evident in more mature Lewy bodies. How this process of mitochondrial degradation influences CHCHD2-specific and overall mitochondrial function is unknown. The relationship of CHCHD2 to its binding partnerCHCHD10 is also of particular interest. Mutation of CHCHD10 can produce ALS/FTD or myopathy, and our proteomics studies also revealed hits associated with AD, ALS and FTD, perhaps suggesting a shared pathophysiologic mechanism between PD and these disorders.

## SUPLEMENTARY MATERIALS

**Supplemental Fig. 1: CHCHD2 T61I mice exhibit normal survival and general locomotor activity through 23 months but increased anxiety. (A)** Survival probability was not different in a second cohort of T61I mice through 23 months of age (Kaplan-Meier survival curve, n = 9 WT, 8 HET and 15 HOM). **(B)** Representative track visualization of homozygous T61I male mice at 23 months (4 WT, 2 HET and 4 HOM) in the open field test. **(C)** Quantifications of total distance and percentage moving time in the open field test using Ethovision. **(D)** Total movement, open arm/total distance, open arm entries and closed arm entries of T61I mice in the elevated plus maze. Female HOM showed trending lower total distance (WT vs HOM *P* = 0.09) and significantly decreased open arm/total distance percentage (WT vs HOM *P* < 0.01, **) and open arm entries (WT vs HOM *P* < 0.01, **) in the elevated plus maze. N = 16 WT, 16 HET and 13 HOM (female), or 14 WT, 18 HET and 17 HOM (male). In all behavioral studies (B-D), data represent mean ± SEM. \*\**P* < 0.01 by linear mixed effects analysis.

**Supplemental Fig. 2: Action potential duration and hyperpolarization-activated current are not altered in CHCHD2 T61I SNc DA neurons. (A)** Spontaneous AP waveforms collected in whole cell current clamp configuration. AP duration was measured from threshold to recrossing threshold during repolarization. AP, action potential. **(B)** *I*_h_ magnitude was measured in whole cell voltage clamp configuration, V_holding_ = −60 stepping to −120 mV. **(C)** Input resistance was measured in whole cell current clamp mode. N = 5 WT and 4 HOM male mice at 23 months. Each circle is a neuron. Data represent kernel density estimations. For all data parametric assumptions were tested to choose between t-test (parametric) or permutation (non-parametric) analysis. \**P* < 0.05. **(D)** Representative images of TH immunoreactivity in striatal sections at 23 months. DA neuron projection areas in dorsal and ventral striatum are indicated with dotted lines. CPu, caudate putamen; NAc, nucleus accumbens; OT, olfactory tubercule. Quantification for TH immunoreactivity showed no difference in WT, HET and HOM by two-way ANOVA with Tukey’s *post hoc* test. N = 4 animals/group. Data represent mean ± SEM. Scale bar indicates 300 μm.

**Supplemental Fig. 3: Increase in T61I CHCHD2 insolubility in the mitochondria but does not impact energy expenditure. (A)** Image of CHCHD2 western blot (left panel). CHCHD2 T61I mutation increased insolubility particularly in the mitochondrial fraction of brain lysate (right panel). N = 2 mice per genotype at 16 months from 2 independent experiments. **(B)** Gel images and quantification of western blot against α-synuclein in Triton-soluble and SDS-soluble fractions of frontal cortex lysate showed no changes in T61I mice at 16 months. A negative control (α-synuclein KO mouse) was included. CB, Coomassie brilliant blue staining as a loading control. α-synuclein level was normalized to total protein content by Coomassie brilliant blue staining. N = 4 WT, 3 HET and 3 HOM. Data represent mean ± SEM. **(C)** Whole-time and dark phase energy expenditure as heat was normal in T61I mice. N = 9 WT, 9 HET and 4 HOM mice at 5 months. Data represents mean ± SEM in bar graphs.

**Supplemental Fig. 4: T61I CHCHD2 VTA DA neurons have decreased expression of Complex I and III genes by Spatial Transcriptomics.** Snap-frozen brain sections of CHCHD2 T61I and KO mice were analyzed by the Visium spatial transcriptomics kit. **(A, B)** Expression of DA neuron markers are confined to spatial regions. A, CHCHD2 T61I; B, CHCHD2 KO. **(C, D)** Volcano plots showing selected metabolic pathways including glycolytic, complex I and complex III genes in the SN (left) and VTA (right) of CHCHD2 T61I (C) and KO (D) mice. **(E)** Bar graphs of glycolytic gene expression of CHCHD2 T61I (up) and CHCHD2 KO (down) mice. The CHCHD2 KO mice showed a significant downregulation of glycolytic genes. **(F)** Bar graphs of complex I component expression of CHCHD2 T61I (up) and CHCHD2 KO (down) mice. CHCHD2 T61I mice exhibited decreased level of nearly all complex I subunits. **(G)** Bar graphs of complex III component expression of CHCHD2 T61I (up) and CHCHD2 KO (down) mice. CHCHD2 T61I mice showed decreased expression of almost all complex III subunits. N = 7 homozygous CHCHD2 T61I mice and 7 littermate controls at 16 months and 4 homozygous CHCHD2 KO and 4 littermate controls at 13 months. In all bar graphs (A, B, E-G), data represent mean ± SEM. \**P* < 0.05, \*\**P* < 0.01 by 2-way ANOVA with Sidak *post hoc* test.

**Supplemental Fig. 5: Distinct mechanisms are implied by individual hits observed in CHCHD2 T61I and KO mice.** Spatial transcriptomics analyses on snap-frozen brain sections of CHCHD2 T61I and KO mice. **(A)** Volcano plots highlighting the top 20 percent individual hits in the SN (left) and VTA (right) of CHCHD2 T61I mice. **(B)** Volcano plots highlighting the individual hits identified in the T61I mice in the SN (left) and VTA (right) of CHCHD2 KO mice. None of the hits was within the top 20 percent in the KO mice. **(C)** Bar graphs of the 3 shared hits in the SN and VTA of the T61I mice, which were unaffected in the CHCHD2 KO mice. N = 7 homozygous CHCHD2 T61I mice and 7 littermate controls at 16 months and 4 homozygous CHCHD2 KO and 4 littermate controls at 13 months. Data represent mean ± SEM in all bar graphs (A, D). \**P* < 0.05, \*\*\**P* < 0.001 by unpaired t-test in D.

**Supplemental Fig. 6: Differentially expressed genes in the CHCHD2 T61I mice are altered in similar trends in cases with Lewy bodies.** Heatmap of normalized gene expression (per region of interest) values from GeoMx spatial transcriptomic analysis of human SNc ventral tier within 8 DEG’s and identified within the CHCHD2 T61I mouse model. N = 10 (59 regions) controls and 5 (27 regions) incidental Lewy bodies (ILB). ‘Limma voom’ methodology was used to assess differential gene expression between control and ILB samples. PMD, post-mortem delay; DV200, RNA integrity number equivalent.

**Supplemental Fig. 7: Assessment of the quality of co-eluted proteins detected from CF-MS experiments. (A)** Agreement of co-eluting proteins from the mitochondrial (mt) and whole cell lysate (WCL) with molecular weights (kDa) inferred from SEC profiles. **(B)** Hierarchical clustering of protein co-fractionation, highlighting normalized peptide count profiles recorded by LC-MS/MS with components of OXPHOS and those linked to PD emphasized. **(C)** auROC curves (sensitivity vs. 1 – specificity) showing the performance of ensemble vs. individual machined learning classifiers such as random forest (RF), generalized linear model (GLM) and support vector machine (SVM) evaluated by fivefold cross-validation using the reference CORUM mouse protein complexes. **(D)** Venn diagram showing the overlap of proteins from each genotype corresponding to the PPIs in Fig. 6C, derived from whole cell lysates and mitochondrial extracts.

**Table S1.** Demographics of the human post-mortem cohorts.

|  | Group | Gender<br>(M/F) | Age at death<br>(years) | Post-mortem delay<br>(hours) |
| --- | --- | --- | --- | --- |
| GeoMx<br>WTA | Control | 3/7 | 92.0 (86.5-94.0) | 25.0 (20.3-34.8) |
|  | ILBD | 3/2 | 88.0 (85.0-95.0) | 20.0 (13.5-35.5) |
|  | ePD | 5/4 | 72.0 (69.0-79.5)<sup>**</sup>, # | 20.0 (6.0-22.5) |
|  | IPD | 5/1 | 77.5 (65.0-80.5)<sup>*</sup> | 15.0 (5.8-18.3) |
| IHC | Control | 3/5 | 25.0 (16.8 -41.8) | 25.0 (16.8-41.8) |
|  | ILBD | 3/2 | 20.0 (13.5-35.5) | 20.0 (13.5-35.5) |
|  | ePD | 2/2 | 19.5 (9.8-27.8)* | 19.5 (9.8-27.8) |
|  | IPD | 6/1 | 15.0 (13.0-22.0) | 15.0 (13.0-22.0) |

Values are presented as median (Q1-Q3). The comparison of age at death among all groups was made with the Kruskal-Wallis test, followed by Dunn's multiple comparisons. The chi-square test was used for gender differences. ILBD = incidental Lewy body disease; ePD = PD cases with early Lewy pathology (Braak stage 4); IPD = PD cases with late Lewy pathology (Braak stage 6). \*, \*\*, compared to controls, $P < 0.05$ and $P < 0.01$ ; #, compared to ILBD, $P < 0.05$ .

## Methods

### Animals

CHCHD2 p.T61I mice were generated using EUCOMM’s conditional-ready embryonic stem (ES) cells. Two single-guide RNAs (5’-GCCTCTGAACCTGGCCCATTTGG-3’ and 5’-TCCTGAACCCTATTTAGTTTTGG-3’) respectively targeting immediate upstream of exon 2 and immediate downstream of exon 3 were used to remove entire exon 2 and 3, and the point mutation was delivered by a donor vector via homologous recombination. ES cells were injected to blastocytes and transferred to recipient females. Mice were maintained on a C57Bl/6 background (The Jackson Laboratory; RRID: IMSR_JAX:000664)

Mice were group-housed in a colony maintained with a standard 12 hours light/dark cycle and given food and water ad libitum. All animal experimental procedures were conducted according to the Guide for the Care and Use of Laboratory Animals, as adopted by the National Institutes of Health, and with approval of the University of California, San Francisco, Institutional Animal Care and Use Committee. Animal care and use in this research are covered under the UCSF ‘‘Assurance of Compliance with PHS Policy on Humane Care and Use of Laboratory Animals by Awardee Institutions’’ number A3400-01.

### qPCR genotyping

CHCHD2 p.T61I genotyping was done by qPCR followed by endpoint allelic discrimination. A product size of 322 bp was amplified using the forward primer (5’-TGAAGATGGCCCAGATCTG-3’) and the reverse primer (5’-CTGAAGCCCCCAGTGAT-3’). Allelic discrimination was done automatically by Applied Biosystem’s SDS 2.4 software using the WT probe (5’ JOE-TTAGATGGCTACCACCGCG-3’ Iowa Black) and the MT probe (5’ 6-FAM-TTAGATGGCGATC ACCGCG-3’ Iowa Black).

### Behavioral testing

General locomotor activity was measured by the open field test using standard procedure(*50*). Mice were placed in the center of a clear acrylic chamber and allowed to explore freely for 15 min. Horizontal and vertical movements were detected using photobeams lining the perimeter of the chamber, and ambulatory and fine movements were separately recorded. Fine movements were defined as breaking the same 2 photobeams repeatedly, while ambulatory movements were defined as breaking 3 or more.

Muscle strength was measured by the inverted grid hang test. Each mouse was first placed with all four paws firmly grasping on a wire grid, which was then inverted so the mouse is hanging over a cage filled with bedding. The latency to fall was recorded, and the maximum latency is 180 s. Mice were tested for 3 trials with an interval of 1 h between each trial.

Motor coordination was measured using a rotarod (Med Associates Inc) as described previously(*51*). Fall from the rotating rod was automatically detected by infrared beams, and the maximum latency was 5 min. 5 mice of the same gender were tested simultaneously in each trial. On the first day, mice were trained to run on a rotating rod at 16 rpm (training phase). Mice were tested on 3 trials with an intertrial interval between 15 and 20 min. On the second and third day of testing, mice were placed on the rod rotating at an accelerating speed from 4 to 40 rpm. The rotation speed increased by 7.2 rpm every 60 s. Mice were tested on 2 sessions of 3 trials with an intertrial interval between 15 and 20 min, and a 2 h interval between the AM and PM sessions.

Anxiety-related locomotion was measured using an elevated plus maze (Kinder Scientific) consisting 2 open arms and 2 closed arms elevated 63 cm above the floor. Mice were given at least 1 h to acclimate to the testing room before mice freely exploring the maze for 10 min. The time, distance, and entries into the open and closed arms were calculated based on infrared photobeam breaks.

### Electrophysiology

Mice were deeply anesthetized with isoflurane and then decapitated. Horizontal brain slices (150 μm) were prepared using a vibratome (Campden Instruments 7000smz-2). Slices were prepared in ice cold Ringer solution (in mM: 119 NaCl, 2.5 KCl, 1.3 MgSO_4_, 1.0 NaH_2_PO_4_, 2.5 CaCl_2_, 26.2 NaHCO_3_, and 11 glucose saturated with 95% O_2_-5% CO_2_) and allowed to recover at 33°C for at least 1 hr. Slices were visualized under an *Axio Examiner A1* equipped with Dodt and IR optics, using a Zeiss Axiocam 506 mono and Neurolucida (MBF Biosciences) software. Whole cell recordings were made at 33°C using 4-5 MΩ pipettes filled with (in mM): 123 K-gluconate, 10 HEPES, 0.2 EGTA, 8 NaCl, 2 MgATP, 0.3 Na_3_GTP, and 0.1% biocytin (pH 7.2, osmolarity adjusted to 275 mOsm/L). Recordings were made using Sutter IPA and SutterPatch software (Sutter Instruments), filtered at 5 kHz and collected at 10 kHz. For all current clamp experiments I = 0. Liquid junction potentials were not corrected during recordings. The reported firing rates are averages of the spontaneous firing rate over the first 2 minutes of whole cell access. *I*_h_ magnitude was measured as the difference between the initial capacitative response to a voltage step from −60 to −120 mV and the final current during the same 800 ms step. Coefficient of Variation was calculated over 100 interspike intervals to evaluate the regularity of firing. Input and series resistances were monitored with hyperpolarizing pulses (0.1 Hz) throughout each experiment. All neurons were filled with biocytin during recording, slices were fixed (4% formaldehyde for 2 hr), and then immunocytochemically processed for TH (*I can add methods if necessary*). Agonists, antagonists, salts, ATP, and GTP were obtained from MilliporeSigma. Quinpirole effects were statistically evaluated in each neuron by binning data into 30 s data points and comparing the last 8 baseline data points to the last 8 data points during drug application. All but 2 neurons recorded in current clamp were quiescent during quinpirole testing; the two firing neurons were excluded from the figure. Both were from WT mice and in both cases quinpirole decreased the spontaneous firing rate.

### Striatal catecholamines analysis

Mice were deeply anesthetized with avertin and perfused with cold phosphate-buffered saline (PBS). Brains were rapidly removed, flash-frozen in a dry ice isopentane bath, and stored at - 80°C before further processing. The striatal region was dissected from half of a brain cut in the sagittal plane, and then trimmed precisely with a cryostat (CryoStar NX50). After the dorsal striatum was fully exposed, punches were made with a pipette tip pre-cut into 1 mm in diameter, and then flash-frozen and stored at −80°C. Catecholamine levels were measured by the Vanderbilt Neurochemistry Core by HPLC coupled to an electrochemical detector (*52*).

### Immunohistochemistry and immunofluorescence staining

Mice were deeply anesthetized with avertin and perfused with cold PBS followed by 4% paraformaldehyde (PFA). Brains were dissected out, postfixed in PFA overnight and stored in 30% sucrose cryoprotectant at 4°C. Brains were cut in the coronal plane into 40 μm-thick sections with a sliding microtome (Leica SM2000R).

For immunofluorescence, brain sections were washed with PBS and transferred in blocking solution containing 0.2% Triton X-100 with 4% BSA for 1 h. Sections were then incubated overnight at 4°C with the appropriate primary antibody (Table S2). Sections were then washed with PBS with 0.2% Triton X-100 and incubated for 2 h at room temperature with the corresponding secondary antibodies (Table S2).

For peroxidase staining, sections were quenched with 3% hydrogen peroxide and 10% methanol in PBS. Sections were then blocked in 10% bovine calf serum with 0.4% Triton X-100. Sections were incubated overnight with rabbit anti-TH (1:1000), mouse anti-α-synuclein (1:500) or rabbit anti-phosphorylated α-synuclein (1:300), followed by biotinylated goat anti-rabbit or anti-mouse IgG (1:300) for 2 h, and then streptavidin-conjugated horseradish peroxidase (HRP) (1:300) for another 2 h. Staining was visualized by developing in 3,3’-diaminobenzidine (DAB, Sigma D12384) and 0.003% hydrogen peroxide solution for 3-5 min.

Brain sections were imaged with a laser scanning confocal microscope (Zeiss LSM880 with Airyscan) or a slide scanner (Leica Aperio Versa). Quantification of fluorescence and area was performed blinded to genotype with ImageJ or CellProfiler.

### Stereology

Total numbers of TH-positive neurons were quantified blinded to genotype. Region selection of SN and VTA was done under 5× magnification following the definition by Fu and colleagues(*53*). Sections were imaged at 63× using a Zeiss Imager microscope (Carl Zeiss Axio Imager A1) equipped with an XYZ computer-controlled motorized stage and an EOS rebel T5i Digital Camera (Canon), and cell counts were performed using MBF Bioscience’s Stereo Investigator. Counting frame size was 60 × 60 μm and systematic random sampling (SRS) grid was 130 × 130 μm, with a section interval of 6.

### Quantifications by Cellprofiler

Acquired images were first segmented to resolve individual cells, by manually drawing each cell using the Fiji rectangle selection tools. Individual cell image files were then saved in the input folder and batch processed in CellProfiler [v4.2.5]. The parameters and modules of our CellProfiler pipelines used in this study can be found https://github.com/szuchiliao/CHCHD2PMMice.git

### α-Synuclein extraction and western blot

Protein was extracted from flash-frozen brains as described by Stojkovska and Mazzuli(*54*). Briefly, homogenized tissues were extracted by agitating in 1% Triton X-100-containing lysis buffer on ice-water slurry for 30 min. 3 freeze-thaw cycles were conducted to facilitate homogenization. Lysate was ultracentrifuged for 30 min at 4°C, and the supernatant was collected as “Triton-soluble fraction”. The pellet was further extracted with SDS lysis buffer, sonicated, and centrifuged for 30 min at 21°C. The supernatant was collected as “SDS-soluble fraction”. Protein concentration was measured using Pierce BCA assay (ThermoFisher, 23227).

15 μg of total protein from the Triton fraction and 25 μg from the SDS fraction were loaded on to polyacrylamide gels (Bio-Rad, 4569033) and electrophoresis was carried out using standard SDS-PAGE procedures. Protein was then transferred from the gel on to a PVDF membrane (Invitrogen, IB24001) using the iBlot^TM^ 2 rapid transfer system (Invitrogen, IB21001). Following transfer, the remaining gel was stained by Coomassie brilliant blue to serve as a loading control. The PVDF membranes were fixed in 0.4% PFA for 30 min at 20°C before blocked for 1 h in a 1:1 mixture of 1X phosphate-buffered saline (PBS) and Li-Cor’s Odyssey PBS blocking buffer (P/N 927-40000). Membrane was incubated overnight with primary antibodies: mouse anti-α-synuclein (BD Transduction Laboratories, 610787, 1:1000), rabbit anti-phosphorylated α-synuclein (phosphor S129; Abcam ab51253, 1:500) and rabbit-anti-Actb (Abcam, ab115777, 1:2500). After incubation in secondary antibodies, Alexa Fluor 647 anti-mouse (Invitrogen, A31571, 1:10000) and Alexa Fluor 488 anti-rabbit (Invitrogen, A11008, 1:10000), membrane was imaged by Bio-Rad’s ChemiDoc Imaging System. Quantification was done by ImageJ.

### Subcellular fractionation

Cells were harvested with a scraper, washed in PBS, homogenized using a glass pestle in isolation medium (250 mM sucrose, 1 mM EDTA, and 10 mM Tris–MOPS, pH 7.4), and then centrifuged at 600 g for 10 minutes at 4°C to pellet nuclei and undisrupted cells. The resulting supernatant was centrifuged at 9000 g for 10 minutes at 4°C, separating the supernatant (containing cytosol) from the pellet. The pellet was resuspended in isolation medium and subjected to 3 additional rounds of centrifugation at 9000 g for 10 minutes at 4°C and the final pellet, containing mitochondria, was retained.

For the isolation of soluble (S) and insoluble (I) fractions from the mitochondrial and cytosolic samples, both fractions were incubated on ice with 1% Triton X-100 for 30 minutes and then centrifuged at 20000 g for 10 minutes at 4°C. After removal of the supernatant (S fraction), the pellet was incubated in SDS buffer for 30 minutes and centrifuged for 10 minutes at 20000 g yielding the insoluble fraction (I fraction).

### Electron Microscopy

Mice were deeply anesthetized with avertin and perfused with cold PBS followed by 4% paraformaldehyde (PFA) and 0.5% glutaraldehyde. Brains were removed and postfixed in the same fixative solution overnight and stored at 4°C. Samples were sectioned into 100 µm thickness with a vibratome. Before immunogold labeling, brain sections were washed with 0.1M phosphate buffer for 3 times, reduced by 1% sodium borohydride for 30 minutes, and incubated in 25% sucrose cryoprotectant for 30 minutes. After 4 more washes, brain sections were flash-frozen with isopentane and 3 freeze-thawed a few cycles to increase permeability. Sections were blocked in 0.1M phosphate buffer with 0.3% BSAc and 0.05% sodium azide, incubated overnight at 4°C with the rabbit anti-TH primary antibody (Millipore AB152) and then with 0.8 nm gold particle goat anti-rabbit antibody (EMS 25100) overnight. Sections were post-fixed for 10-minutes in 1.5% glutaraldehyde solution at room temperature. After 2 washes with 0.1M phosphate buffer and 5 washes with ddH_2_O, silver enhancement was done for 7 minutes at room temperature (Nanoprobes, 2012).

### Post-mortem human substantia nigra

Tissue samples from pathologically confirmed cases with incidental Lewy bodies (ILB, n=5), early PD (n=9), and late PD (n=8), and controls without the neurological or neuropathological disease (n=10) were obtained from the Sydney Brain Bank (Table S1). The study was approved by the University of Sydney Human Research Ethics Committee (2021/845). All cases with PD were levodopa-responsive and fulfilled the UK Brain Bank Clinical Criteria for the clinical diagnosis (*55*) with no other neurodegenerative conditions. Controls had no Lewy pathology, while ILB and PD cases were staged using the regional location of the Lewy pathology (*56, 57*). Formalin-fixed paraffin-embedded (FFPE) sections from controls, ILB (Braak stage I-II), early PD (Braak stage IV), and late PD (Braak stage VI) were used to examine the expression of *CHCHD2*, *CHCHD10*, and the locational of their proteins compared with intracellular αSyn aggregations in DA neurons.

### Human Tissue preparation and immunofluorescence staining

FFPE blocks of human midbrains containing substantia nigra (SN) were cut on a rotary microtome (HistoCore MULTICUT R Rotary, Leica Biosystems) at 6 µm. Sections were mounted on Fisher Scientific (12-550-15) for GeoMx Human Whole Transcriptome Atlas (WTA, nanostring) according to the manufacturer’s instruction or on Series 2 adhesive microscope slides (Trajan Scientific Medical, AU) for immunofluorescence. Before staining, sections were de-waxed and rehydrated with HistoChoice® Clearing Agent (Sigma, H2779) and gradient concentrations of ethanol. Each antibody was tested with immunohistochemistry to decide the optional solution for heat-induced antigen retrieval. For this immunofluorescence approach, formic acid (70% for 20 mins) was applied, followed by Tris-EDTA (pH 9.0) in a programmable antigen retrieval cooker (Aptum Bio Retriever 2100, Aptum Biologics Ltd, UK). Sections were blocked with Human BD Fc Block (BD Pharmingen, 564219, 1:50) for 30 mins at room temperature, then TrueBlack® Background Suppressor (Biotium, 23012A) for 1 hr at RT. Primary antibodies (either CHCHD2 or CHCHD10 with αSyn and mitochondrial markers) were diluted in blocking buffer (2% donkey serum, 1% BSA, 0.1% fish gelatine, 0.3% Triton X-100, 0.1% Tween-20, and 0.02% NaN_3_ in PBS) for section incubation at RT for one overnight, followed by the corresponding Alexa Fluor secondary antibodies with Hoechst nucleus counterstaining (bisBenzimide H 33342 trihydrochloride, Sigma, B2261, 1 μg/ml) at RT for 2 hrs. Sections were further incubated in Alexa flour 790 conjugated TH antibody in blocking buffer at RT for 2 hrs before being treated with TrueBlack® Lipofuscin Autofluorescence Quencher (Biotium, 23007). Slides were mounted with EverBrite™ mounting medium (Biotium, 23001) and sealed with Biotium’s CoverGrip™ Coverslip Sealant (23005). Negative controls were performed for each staining batch, omitting primary or secondary antibodies.

### Metabolomics

Mice were fasted overnight prior to anesthetizing with isoflurane. The mice then received a 200 μl bolus of [13-C]glucose solution (Cambridge Isotope Labs, CLM-1396-1) via intraperitoneal injection, followed by tail vein infusion at a rate of 150 μl/h. After infusion, mice were decapitated and the brains rapidly removed and flash-frozen in dry-iced isopentane. Metabolites were extracted from homogenized brains by incubation in 80% methanol at −80°C for 20 min, followed by centrifugation at 14,000 g for 15 min. Metabolite supernatants were dried in a Labconco CentriVap prior to storage at −80C. Samples were quantified at the UCLA Metabolomics Core.

### Spatial transcriptomics

#### Mouse

Spatial transcriptomics were acquired with Visium spatial gene expression kits (10X Genomics). Sample preparation, sample imaging, and library generation were completed in accordance with 10X Spatial Gene Expression protocols. Briefly, fresh brain tissue was flash frozen in a dry ice isopentane bath. The brain tissue was then embedded in Optimal Cutting Temperature (OCT) compound (Tissue-Tek 62550-12). 10 µm-thin section from the midbrain was made with a cryostat (CryoStar NX50) and then mounted onto a 10X spatial gene expression slide. Sections were stained with TH, NeuN, and Hoechst 33342 before imaging on a Leica Aperio Versa slide scanner. The cDNA libraries were generated at the Gladstone Genomics Core and sequenced at the UCSF Center for Advanced Technology on an Illumina NovaSeq 600 on an SP flow cell. Alignment of the sequencing data with spatial data from the Visium slide was completed with the 10X Space Ranger software. Two anatomical regions of interest, SN and VTA, and a control region, thalamus, were identified. RNA capture areas corresponding to each anatomical region were selected for analysis based on their spatial proximity to the anatomical regions and on the expression of known genetic markers. SN and VTA genetic markers included TH, DAT, and VMAT2; thalamus markers included Prkcd, Ptpn3, and Synpo2. Demarcation of SN and VTA was done according to Fu et al., 2012. SN and VTA capture areas also had to contain at least one complete DA neuron soma. Gene rankings for hit analysis were established using the fold change score (FCS) and signal-to-noise score (SNS). Equations for these scores are given as:

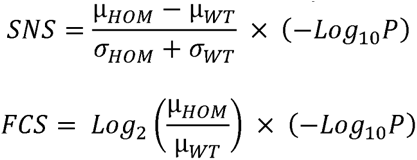

where µ is the average gene expression, σ is the standard deviation, and P is the p-value derived from a t-test. Genes with a p-value < 0.05 that also appeared in the top 20% for both scoring metrics were highlighted as differentially expressed genes of interest.

#### Human

FFPE blocks of post-mortem human midbrains were sectioned at 6 µm thickness using a rotary microtome (HistoCore MULTICUT R Rotary, Leica Biosystems). The sections were mounted on Fisher Scientific slides (catalog #12-550-15) and processed using the Nanostring GeoMx® Digital Spatial Profiler according to the manufacturer’s instructions, isolating transcripts from either full regions of interest (unmasked) or TH+ (masked) regions (∼500 µm). Sequencing libraries were constructed using the Human Whole Transcriptome Atlas (GeoMx Hu WTA) and sequenced on the NovaSeq 6000 following manufacturer protocols. Technical replicates were merged using the Linux ‘cat’ command, and data alignment and feature counting were conducted via the GeoMx NGS analysis pipeline (version 2.0.21) on the Illumina BaseSpace platform. Quality control procedures were executed in R statistical software, applying the following thresholds: minimum segment reads of 1000, ≥80% for percent trimmed, stitched, and aligned, ≥50% for percent saturation, a minimum negative count of 1, a maximum NTC count of 10,000, a minimum of 20 nuclei, and a minimum area of 1000. The overall gene detection rate across all samples was set at a minimum of 1%, with a per-sample minimum gene detection rate also set at 1%. The sequencing of retrieved mRNA probes facilitated spatial transcriptomic (ST) measurements of 18,675 transcripts, of which 9,012 passed the limit of quantification. Raw counts were subsequently normalized using quantile normalization and DEGs evaluated using LIMMA Voom.

### Code availability

Scripts used for low level Nanostring GeoMx data processing are available https://github.com/zchatt/ASAP-SpatialTranscriptomics/tree/main/geomx

Scripts used for higher level data analysis (DEG, heatmap etc.) are available https://github.com/zchatt/ASAP-SpatialTranscriptomics/tree/main/Liao_2024

### Co-fractionation coupled with mass spectrometry (CF-MS)

Whole cell extracts (WCE) and mitochondrial extracts (mt) totaling 500 µg each were obtained from the brains of CHCHD2 WT, CHCHD2 T61 heterozygous and homozygous, and CHCHD2 KO mice. These samples were cross-linked using dithiobis (succinimidyl propionate) (DSP) and then subjected to size exclusion chromatography (SEC) employing an Agilent 1100 HPLC system equipped with a 300 × 7.8 mm BioSep-4000 Column (Phenomenex), in accordance with a protocol recently described in the literature (*39*). This procedure yielded 96 fractions for each extract per genotype. These fractions were subsequently digested with trypsin and analyzed via an Easy-nanoflow liquid chromatography 1200 (Easy nLC; Proxeon) system connected to an Orbitrap Exploris mass spectrometer (Thermo Fisher Scientific). The method for processing the digested samples, the chromatographic separation, and the settings for the comprehensive scanning of mass spectrometry spectra are detailed as previously (*39*).

### Liquid chromatography tandem mass spectrometry (LC-MS/MS)

NanoLC connected to an Orbitrap Exploris mass spectrometer (Thermo Fisher Scientific) was used for the analysis of all samples. The peptide separation was carried out using a Proxeon EASY nLC 1200 System (Thermo Fisher Scientific) fitted with a custom-made C18 column (15 cm x 150 μm ID) packed with HxSil C18 3 μm Resin 100 Å (Hamilton). A gradient of water/acetonitrile/0.1% formic acid was employed for chromatography. The samples were injected onto the column and run for 95 minutes at a flow rate of 0.60 μl/min. The peptide separation began with 1% acetonitrile, increasing to 3% in the first 2 minutes, followed by a linear gradient from 3% to 23% acetonitrile over 74 minutes, then a steep increase from 24% to 80% acetonitrile over 10 minutes, and finally a 10-minute wash at 80% acetonitrile. The eluted peptides were ionized using positive nanoelectrospray ionization (NSI) and directly introduced into the mass spectrometer with an ion source temperature set at 250°C and an ion spray voltage of 2.1 kV. Full-scan MS spectra (m/z 350–2000) were captured in Orbitrap Exploris at a resolution of 240,000 (m/z 400). The automatic gain control was set to 1e6 for full FTMS scans and 5e4 for MS/MS scans. Ions with intensities above 1500 counts underwent fragmentation in the linear ion trap. The top 15 most intense ions with charge states of ≥2 was sequentially isolated and fragmented using normalized collision energy of 30%, activation Q of 0.250, and an activation time of 10 ms. Ions selected for MS/MS were excluded from further selection for 30 seconds. The Orbitrap Exploris mass spectrometer was operated using Thermo XCalibur software.

### LC-MS/MS analysis

Raw spectral files from IP-MS or size exclusion chromatography-high performance liquid chromatography (SEC-HPLC) fractionation-MS experiments were converted into mzXML format using ReAdW software and then searched against the canonical mouse protein sequences from the UniProt database. To enhance the sensitivity and accuracy of peptide identification, the mzXML files underwent searches against these mouse protein sequences utilizing peptide-spectrum matches from three distinct search engines: Tide (from the Crux suite version 4.1), MS-GF+ (version 2017.01.13), and MaxQuant (version 1.6.7.0) for CF-MS, while Tide and MS-GF+ for IP-MS experiments. These searches aimed to maintain a false discovery rate (FDR) below 0.1% for Tide and MS-GF+ or 1% for MaxQuant, both for peptide and protein identifications. The results from Tide and MS-GF+ were further refined using Percolator1. MaxQuant searches were conducted with a 4.5 ppm tolerance for precursor mass in the main search, a 20 ppm tolerance for fragment ion or peptide mass in the first search, and allowance for up to two missed cleavages. Tide searches were set with a 10 ppm tolerance for fragment ion mass and one missed cleavage, while MS-GF+ searches allowed a 20 ppm tolerance for precursor mass with a percolator peptide Q value threshold set at 0.1. Across all searches, carbamidomethylation of cysteine was treated as a fixed modification, with methionine oxidation and protein N-terminal acetylation permitted as variable modifications.

### Computational scoring to predict protein-protein interactions (PPI) from CF-MS datasets

For predicting endogenous protein interactions from CF-MS experiments using whole cell lysate (WCL) and mitochondrial (mt) extracts, the MACP algorithm (PMID: 37985878) was employed. The preprocessing steps undertaken included: (1) Averaging Log2-transformed MaxQuant MS1 ion intensities together with spectral counts from Tide and MS-GF+; (2) Excluding proteins identified in only one fraction; (3) Imputing zeros for missing proteins not detected by MS, due to either low abundance or absence in the mouse tissues; and (4) Adjusting for fraction bias and discrepancies in sample injection through row- and column-wise normalization, accounting for the variation in peptide abundance across fractions. Subsequently, the protein elution profile matrix from each search engine for a given sample was independently analyzed for PPI scoring. This analysis employed 17 distinct correlation metrics to calculate pairwise protein profile similarities, with details of each profile feature have been described in our recent study(*39*). The results from each search engine were then integrated using a random forest (RF) classifier, which had been trained using PPIs from curated mouse complexes in the CORUM^4^ and IntAct^5^ databases. Scores for each protein pair across different search engines were merged to compute an average RF score. The model’s prediction performance was evaluated through 10-fold cross-validation against an independent set of PPIs, establishing an RF cut-off score of > 0.5. Protein pairs scoring below this 0.5 threshold were excluded from consideration.

### Multimeric protein complex predictions from PPI network

High-confidence PPIs identified through CF-MS experiments across different genotypes were analyzed using the ClusterONE (Clustering with Overlapping Neighborhood Expansion) algorithm to delineate complex memberships. The optimization of ClusterONE involved various combinations of parameter adjustments (density, d; overlap, o) alongside a suite of evaluation metrics (accuracy, maximum matching ratio, overlap) as we previously described (*39*). In brief, the predicted complexes were compared against known CORUM and IntAct mouse protein complexes using a range of density (0.2-0.45) and overlap (0.5-0.8) settings across the evaluation metrics. The parameter settings that yielded the highest composite score— representing the sum of accuracy, overlap, and maximum matching ratio—were deemed most effective for identifying high-quality protein complexes. The interactions among proteins across the four genotypes, along with the predicted macromolecular complexes from the PPI networks derived from CF-MS datasets, were visualized using Cytoscape^7^ software (ver. 3.9.1).

### Differential protein complex analyses

To assess the complex rewiring among the four genotypes, we utilized coelution profiles of the predicted protein complex subunits. Gene set enrichment analysis (GSEA) was performed, treating each complex identified within the four genotypes as a gene set. We then compared the coelution profiles of each subunit within these gene sets across the genotypes.

### Complex I activity assay

A detailed protocol for characterizing isolated mitochondria can be found at https://doi.org/10.17504/protocols.io.e6nvwdz87lmk/v1. A piece of mouse cerebrum containing the midbrain was flash-frozen and pulverized for mitochondrial isolation. The mass of isolated mitochondria was measured using a Pierce BCA protein assay kit. To measure enzyme activity of complex I, 10 μg of crude mitochondria preps was dispensed into a 96-well plate containing 50 mM pH 7.4 potassium phosphate buffer, 2 mM potassium cyanide, 2.5 mg/mL bovine serum albumin, 5 mM MgCl2, 2 μM Antimycin A, 100 μM decylubiquinone, and 300 μM K2NADH, with or without 1 μM rotenone. Immediately after combining mitochondria with the above reagents, 340 nm absorbance was measured for 45 min on a Spectramax M4 plate reader.

### Key resources table

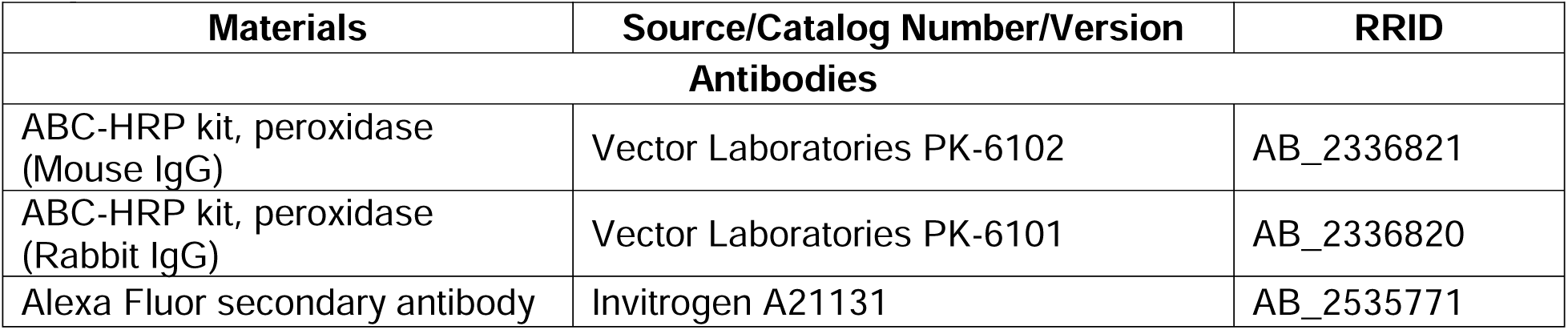
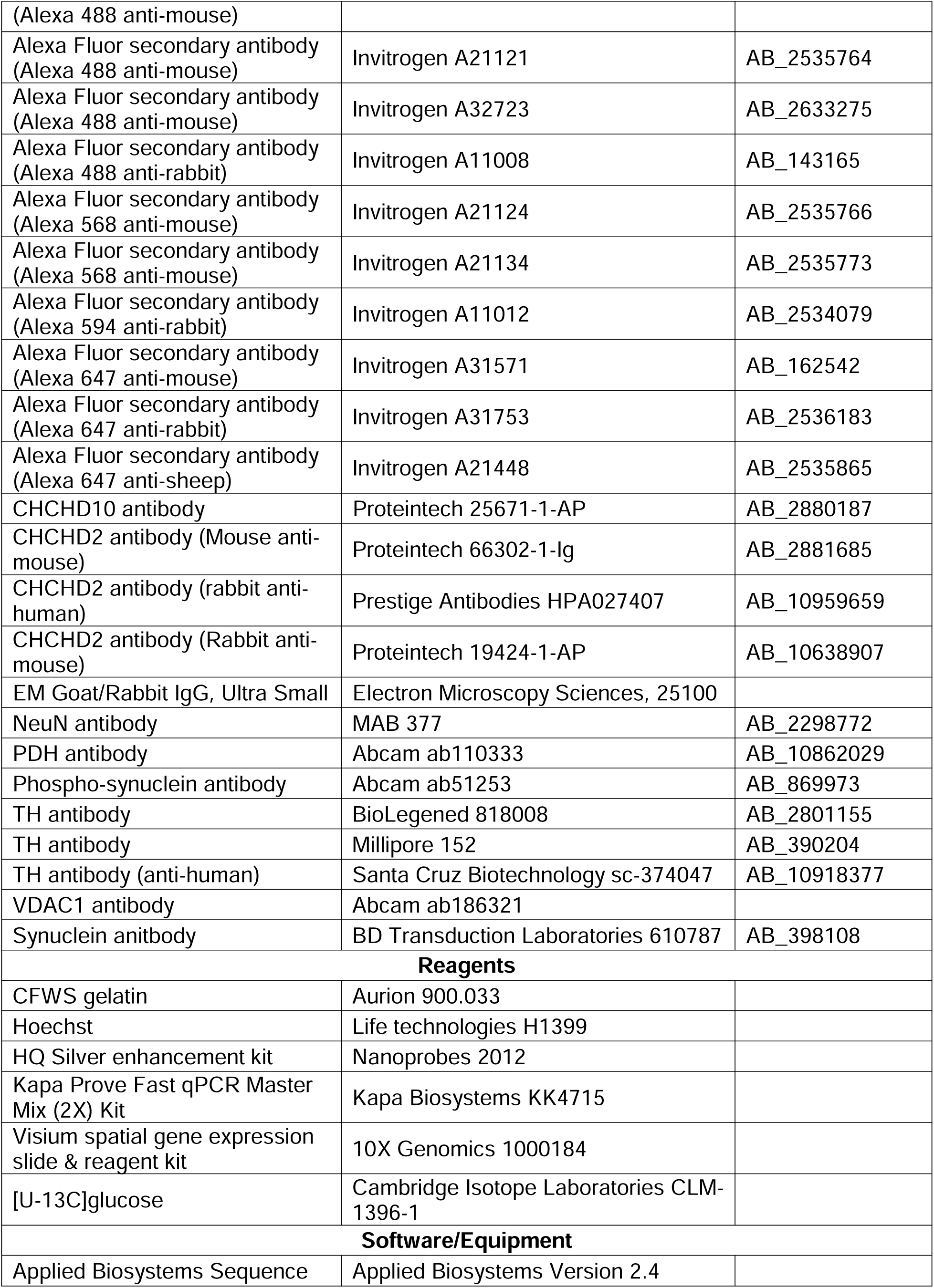
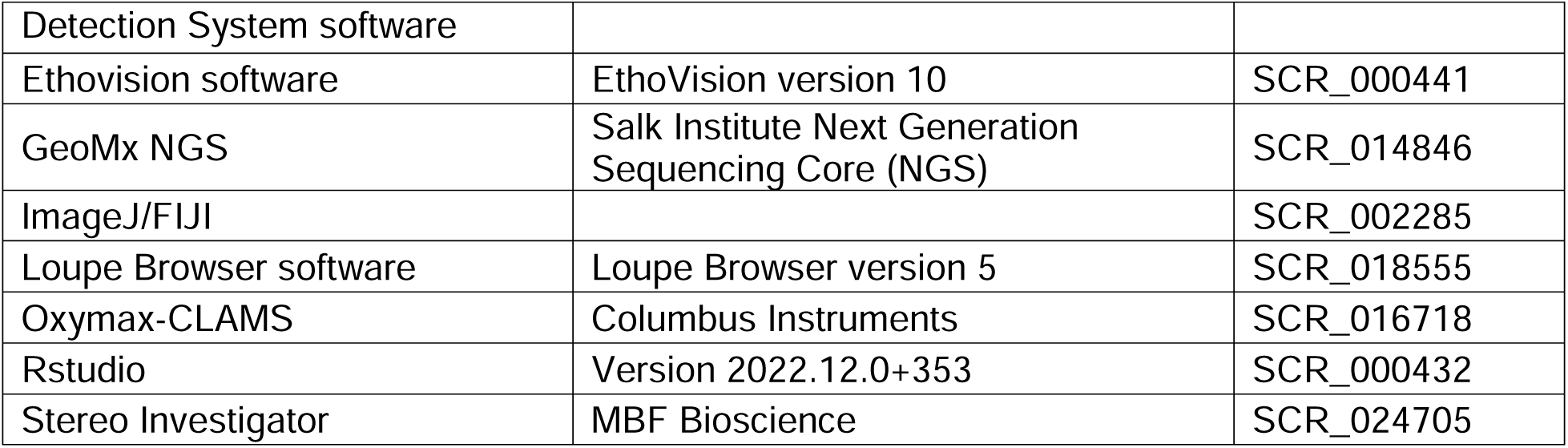

## Acknowledgments

We thank Jessica Speckart, Julia Holtzman, and Iris Lo in the Gladstone Behavioral core for their assistance with behavioral testing and data analysis, Hongyun Li for antibody testing and Ping Wu for the human brain tissue staining, and Johanna ten Hoeve, Thomas G. Graeber and the UCLA Metabolomics Center for their assistance with metabolomics studies and data processing. We also thank the Gladstone Histology and Light Microscopy core, as well as Kathryn Claiborn for helping edit the manuscript, Tami Tolpa for help with graphics, and Erica Delin for administrative assistance.

