## Supplementary Figures for "CHCHD2 mutant mice display mitochondrial protein accumulation and disrupted energy metabolism"

### Supplementary Figure 1

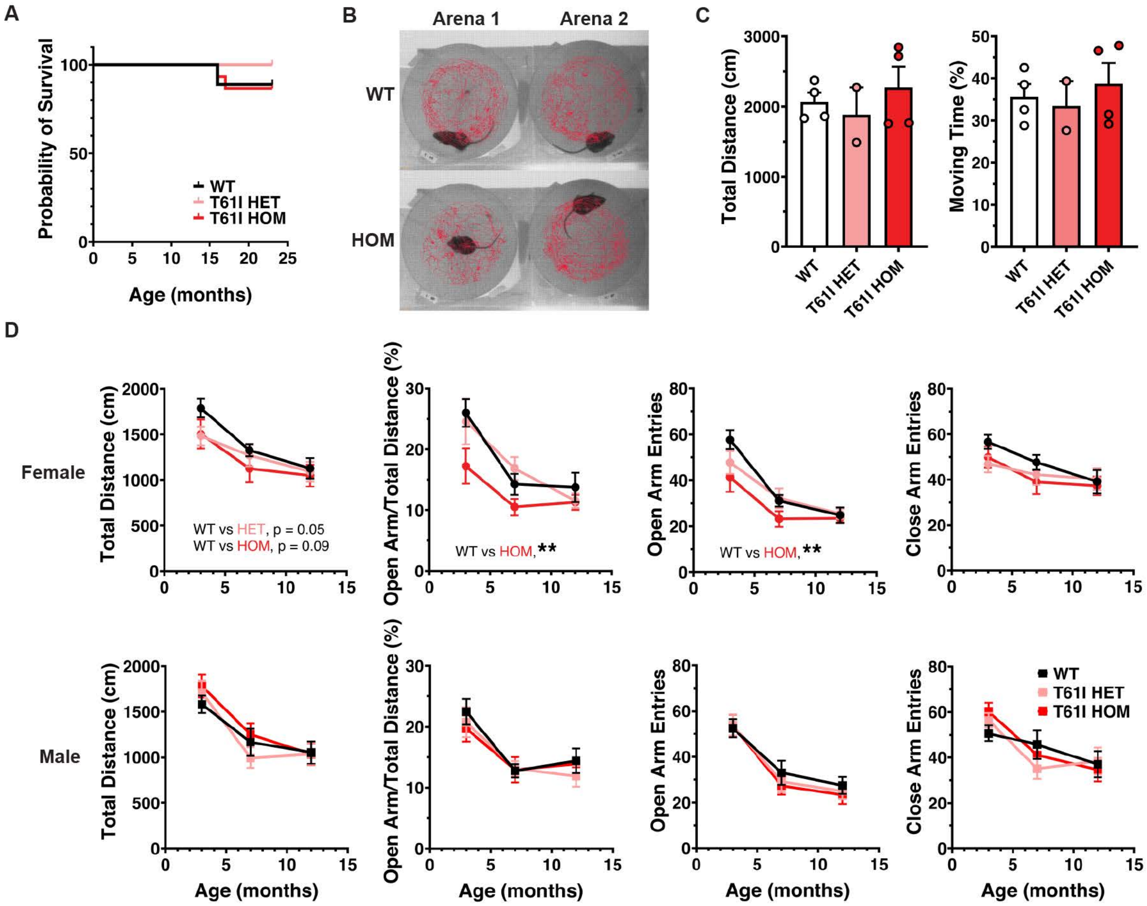

Supplementary Figure 2

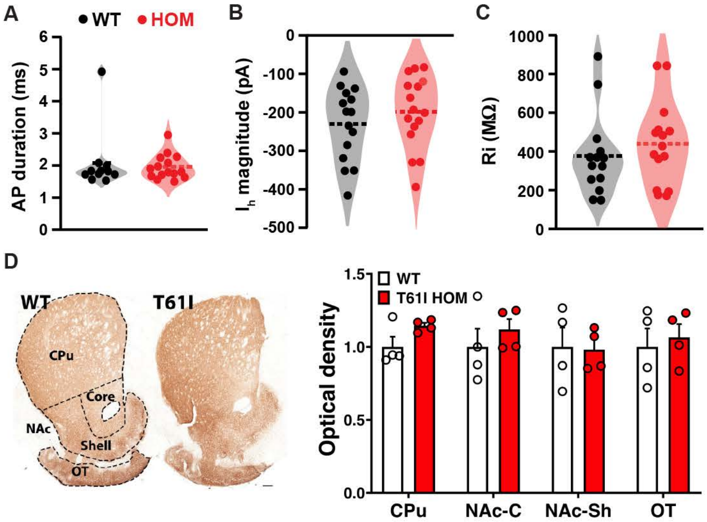

Supplementary Figure 3

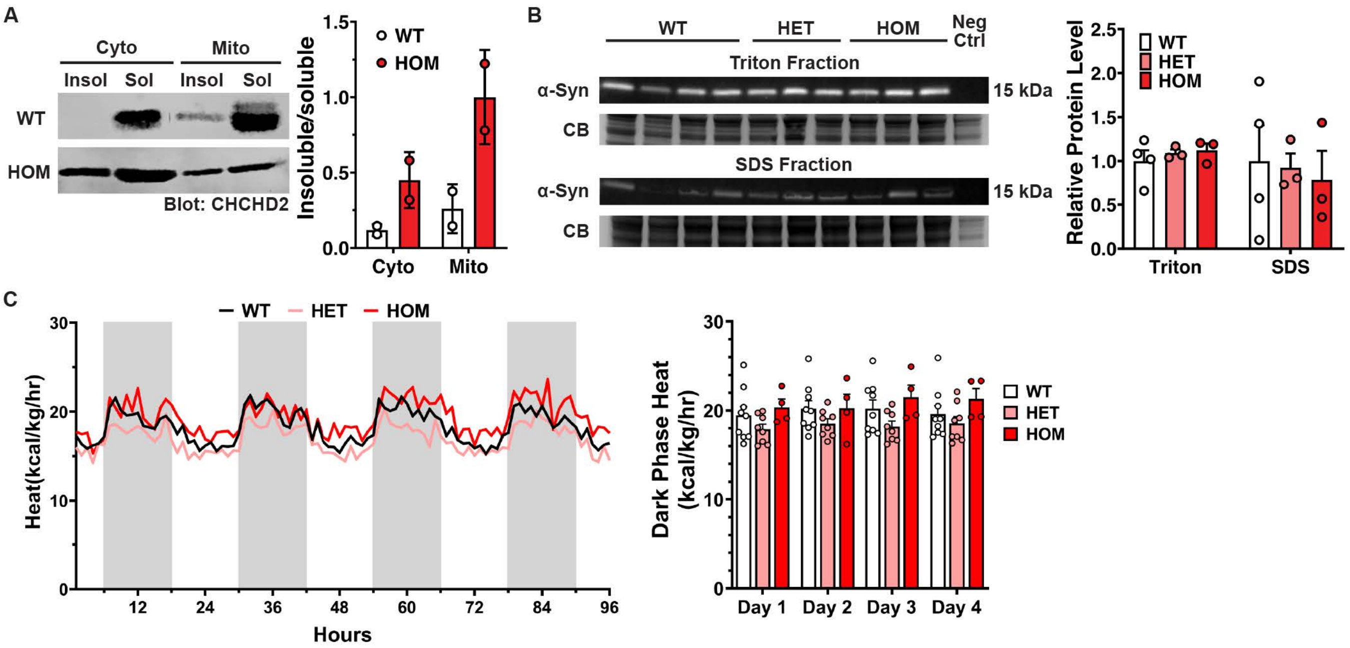

**Supplementary Figure 4**

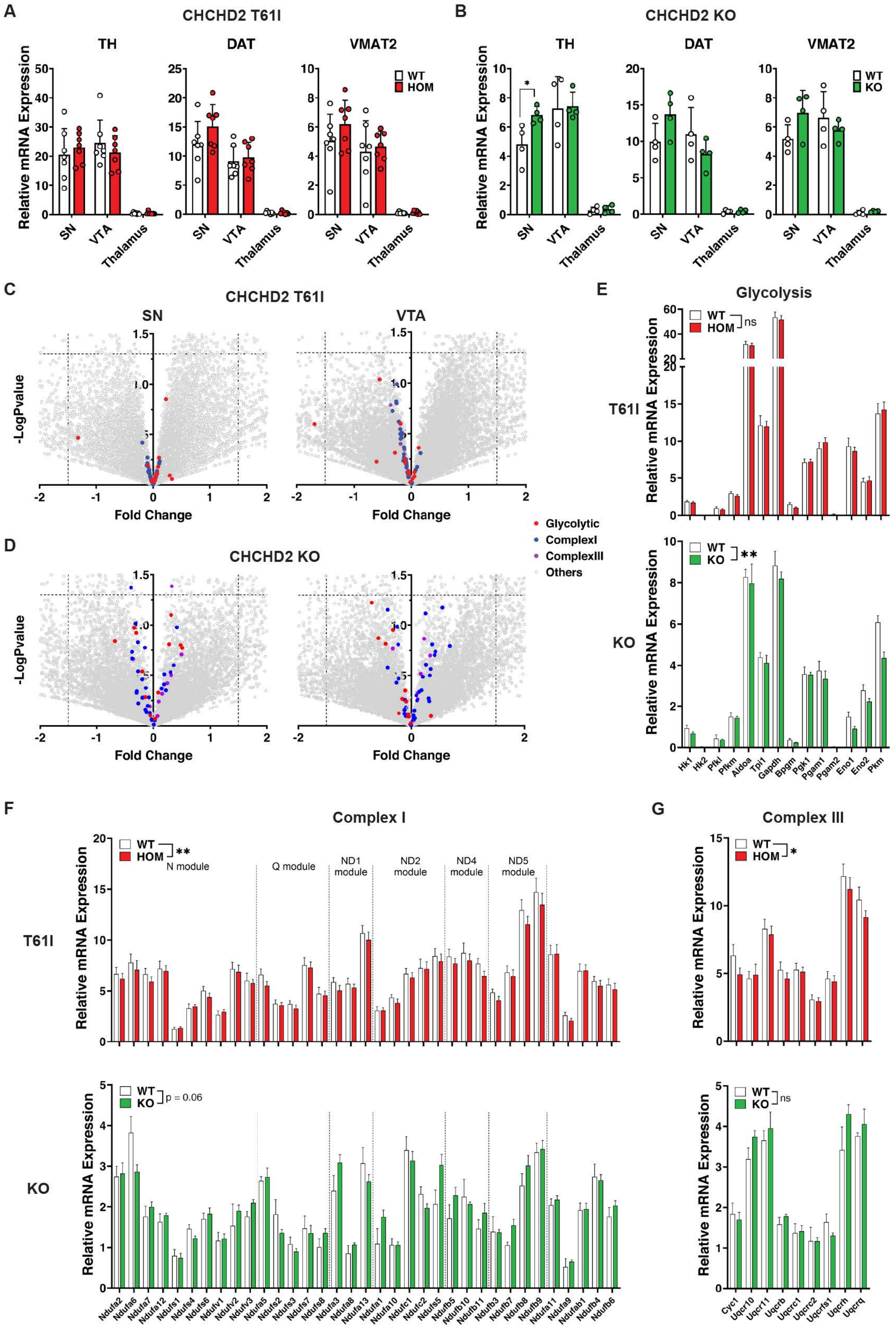

### Supplementary Figure 5

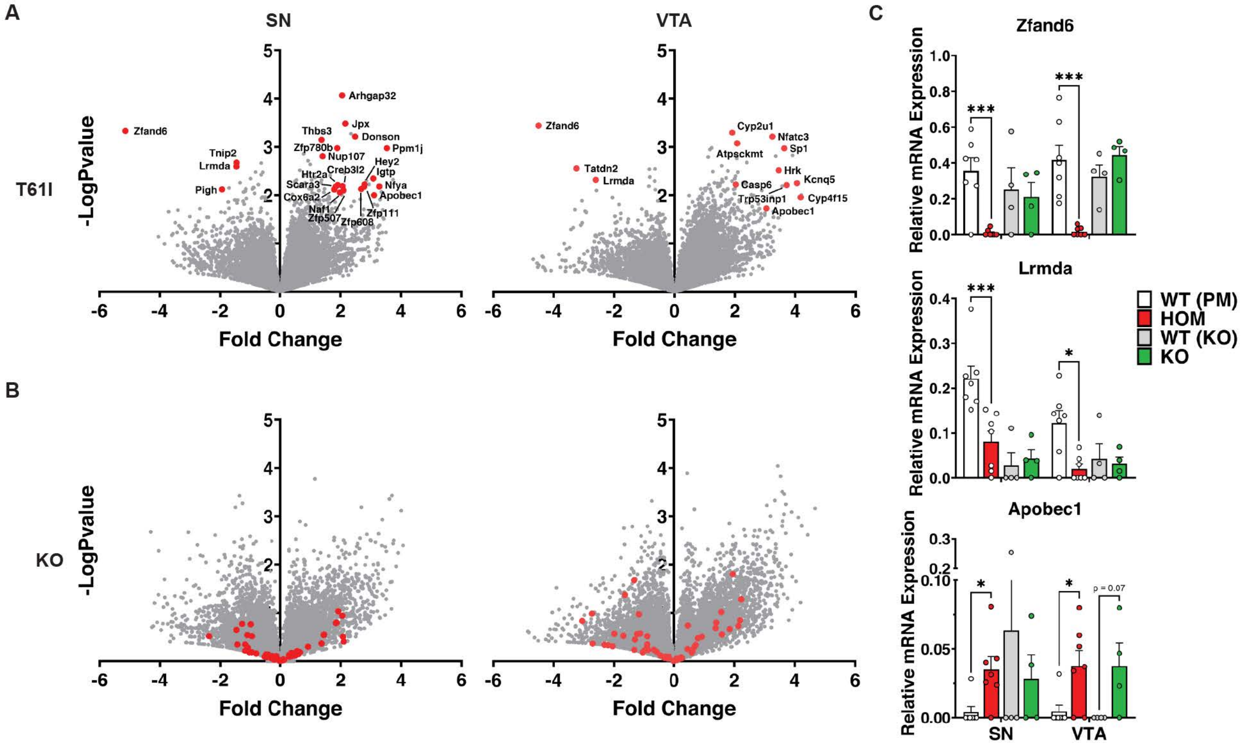

**Supplementary Figure 6**

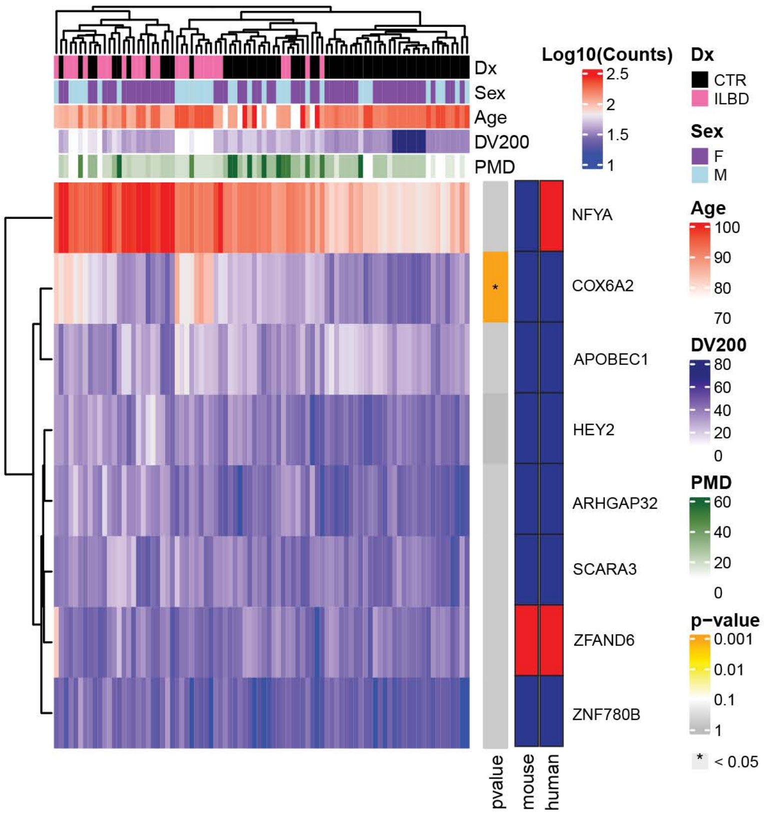

### Supplementary Figure 7

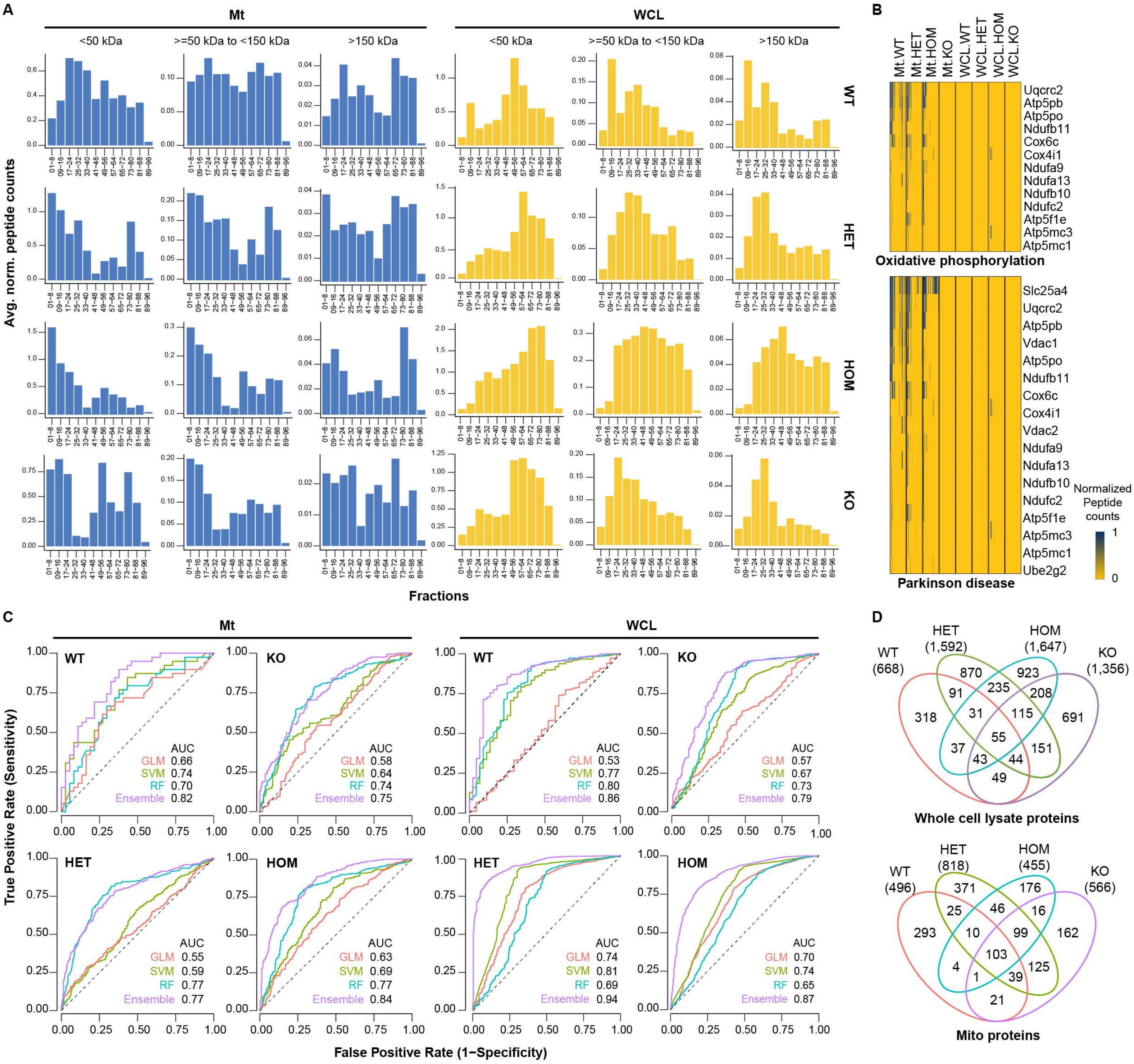
